# Enhanced Hippocampal Spare Capacity in Q175DN Mice Despite Elevated mHTT Aggregation

**DOI:** 10.1101/2024.10.14.618355

**Authors:** Melissa A Solem, Ross Pelzel, Nicholas B. Rozema, Taylor G. Brown, Emma Reid, Rachel H. Mansky, R Gomez-Pastor

## Abstract

**Background:** Huntington’s disease (HD) is a neurodegenerative disease resulting in devastating motor, cognitive, and psychiatric deficits. The striatum is a brain region that controls movement and some forms of cognition and is most significantly impacted in HD. However, despite well-documented deficits in learning and memory in HD, knowledge of the potential implication of other brain regions such as the hippocampus remains limited.

**Objective:** Here, we study the comparative impact of enhanced mHTT aggregation and neuropathology in the striatum and hippocampus of two HD mouse models.

**Methods:** We utilized the zQ175 as a control HD mouse model and the Q175DN mice lacking the PGK-Neomycin cassette generated in house. We performed a comparative characterization of the neuropathology between zQ175 and Q175DN mice in the striatum and the hippocampus by assessing HTT aggregation, neuronal and glial pathology, chaperone expression, and synaptic density.

**Results:** We showed that Q175DN mice presented enhanced mHTT aggregation in both striatum and hippocampus compared to zQ175. Striatal neurons showed a greater susceptibility to enhanced accumulation of mHTT than hippocampal neurons in Q175DN despite high levels of mHTT in both regions. Contrary to the pathology seen in the striatum, Q175DN hippocampus presented enhanced spare capacity showing increased synaptic density, decreased Iba1^+^ microglia density and enhanced HSF1 levels in specific subregions of the hippocampus compared to zQ175.

**Conclusions:** Q175DN mice are a valuable tool to understand the fundamental susceptibility differences to mHTT toxicity between striatal neurons and other neuronal subtypes. Furthermore, our findings also suggest that cognitive deficits observed in HD animals might arise from either striatum dysfunction or other regions involved in cognitive processes but not from hippocampal degeneration.

## INTRODUCTION

Huntington’s disease (HD) is an inherited neurodegenerative disease caused by a poly-glutamine expansion in exon 1 of the *HTT* gene resulting in misfolding and aggregation of the mutant HTT protein (mHTT) [1–3]. The striatum is most significantly impacted in HD and is the focus of most HD studies [4]. Despite mHTT aggregation being observed across the brain, the effects of its toxicity demonstrate a high degree of cell-specificity, with medium spiny neurons (MSNs) of the striatum being selectively vulnerable to mHTT[5–10]. However, the mechanisms that lead to the enhanced MSN susceptibility in HD are not yet resolved.

HD is primarily characterized as a motor disease, but it also manifests with cognitive and behavioral deficits that appear well before motor symptoms onset and are considered the most burdensome for patients and caregivers [11–15]. The earliest and most sensitive indicators of HD include impairment of executive functions, difficulty learning new information and retrieving previously learned information, and changes to cognitive processing and cognitive flexibility (CF) [13]. Deficits in declarative memory, involving memories of facts and events [16], are also widely reported [12, 17–20]. Despite the progress made in characterizing the different cognitive aspects of HD and developing reliable measures of cognition in prodromal and other stages of the disease, very little is known about the brain regions and mechanisms involved in the onset and progression of cognitive symptoms in HD. Although emerging evidence reflects an involvement of the striatum in several forms of cognition [21–26], the hippocampus remains widely considered a structure critical to learning and memory [27, 28]. Unfortunately, knowledge of the potential implication of the hippocampus in HD-related cognitive deficits remains limited due to the sparse studies in HD models addressing this problem [29–31]. Recent cognitive testing and neuroimaging studies of patients indicate that hippocampal dysfunction and degeneration may arise in the early to mid HD, the stage associated with the emergence of cognitive and psychiatric symptoms [32–35]. However, definitive analyses involving the degeneration of the hippocampus in cognitive decline in HD mice are lacking.

The full-length knock-in zQ175 model (C57BL/6 background) has recently become one of the most commonly used mouse models in HD research. zQ175 mice express a chimeric mouse/human exon 1 of *Htt* containing ∼188 CAG repeats under the endogenous mouse *Htt* promoter [36]. This mouse model recapitulates MSN neurodegeneration, striatal neuropathology, as well as several behavioral phenotypes observed in HD patients. However, although symptoms onset occurs earlier than in other well established HD knock-in models such as the Q111 and Q140 [37, 38], it is still deemed as a mild progressive model with only subtle behavioral deficits arising between 3-4 months of age. A major caveat of this model, but also other models of HD, is the lack of overt MSN loss as it is observed in HD patients [39–42]. Studies of hippocampal pathology in the zQ175 mouse model are scarce. HTT aggregates have been reported in the hippocampus of zQ175 mice, especially with antibodies targeting HTT exon 1, although the level of aggregation is significantly lower than in other parts of the brain [30]. A recent study using small-animal PET analyses with different radioligands showed a modest binding decline in D1 and 5-HT2A receptors in the hippocampus of zQ175 mice compared to WT [43], suggesting some hippocampal alterations, although the functional implication of these findings has not been explored.

Additional efforts have been made to create better and more robust HD mouse models [44, 45]. One example is the zQ175 mouse backcrossed on the FVB/N background (Q175F) lacking the Neomycin cassette present upstream of the *Htt* gene (Q175FDN)[45]. Southwell et al. showed Q175FDN mice enhanced mHTT expression levels and aggregation compared to the Q175F and presented an earlier onset of HD-like motor and cognitive phenotypes [45]. Hippocampal synaptic dysfunction has been reported in Q175FDN [46, 47] although these studies did not include comparisons with the parental Q175F line due to the Htt deficiency phenotypes observed in the neo intact line and sudden death from fatal seizures, and therefore it is hard to determine if such effects were present or not in the parental line. However, in the characterization of the Q175FDN line, heterozygous and homozygous genotypes were compared showing a more severe phenotype in homozygous mice. A caveat of the Q175FDN is the severe visual impairment characteristic of the FVB background [48] that makes behavioral experimentation requiring visual cues difficult [49], which is often implemented in tasks assessing cognitive functions. To alleviate these caveats, the CHDI in collaboration with the Jackson Laboratory generated the Q175DN using the parental zQ175 mouse line in the C57BL/6 background, which has been the subject of recent in-depth characterization and the preferred model to assess recent disease biomarkers[50–53]. Others have independently generated the Q175DN line reproducing the enhanced pathology and symptomatology of this mouse line [54].

In our study we have investigated the suitability of the heterozygous Q175DN mouse line (generated in house) in the study of hippocampal pathology by direct comparison with the heterozygous zQ175 parental line. Our results demonstrate that Q175DN mice exhibit significantly enhanced mHTT aggregation in the striatum and enhanced striatal neuropathology, shown by decreased expression of striatal neuron markers compared to zQ175, as previously reported [45]. Importantly, Q175DN mice also exhibited significantly greater levels of HTT aggregation in the hippocampus, compared to zQ175 mice. Intriguingly, despite increased mHTT aggregation in the hippocampus, no markers of hippocampal pathology were observed through analyses investigating neuroinflammation, cortical layer thinning, chaperone expression, and synaptic density. More importantly, we found that while striatal pathology worsened in Q175DN mice, some regions of the hippocampus displayed enhanced synaptic density, decreased Iba1^+^ cells, and increased HSF1 levels. Our findings suggest an enhanced spare capacity of hippocampal neurons to mHTT toxicity and imply that cognitive deficits reported in Q175DN might not directly result from overt hippocampal degeneration. Our data further demonstrates the selective vulnerability of striatal neurons in HD and suggests that the Q175DN mouse model may serve as a valuable tool to understand the fundamental susceptibility differences to mHTT toxicity between different neuronal subtypes.

## MATERIALS AND METHODS

### Mouse Lines

For this study, we used the heterozygous zQ175 on the C57BL/6 background lacking the neomycin cassette, referred hereafter as Q175DN. These mice were generated when the parental heterozygous zQ175 line (referred hereafter as zQ175 for the presence of Neomycin) was crossed with the Gpr88-Cre line (Stock #022510)[55] (**Supplementary Figure 1**). Although the *Gpr88* promoter was designed to specifically express Cre in MSNs [56], there was a spontaneous expression in the germline in a fraction of the mouse crosses causing a systemic deletion of the Neo cassette. Once the deletion was identified in tail genotyping, Q175DN mice were selected and propagated. Genotyping of zQ175 was conducted with primers Hdh-I: 5’-CATTCATTGCCTTGCTGCTAA-3’, Hdh-II: 5’-CTGAAACGACTTGAGCGACTC-3’, Hdh-III: 5’-AAAGGAAAAGGAAAAAGCCAAGC-3’, Hdh-IV: 5’-TCCCTGACTAGGGGAGGAGTAGAA-3’. Genotyping of Q175DN was conducted with primers DNeo-I: 5’-GCGGGCTTATACCCCTACAG-3’, DNeo-II: 5’-TCCAGGACAGCCAGAGCTAC-3’. WT (C57BL/6) animals were used as controls. Animals were used between 8 and 15 months of age (WT; ∼10-13mths, zQ175; ∼10-14 mths with one mouse being 8 mths, Q175DN; ∼10-15 mths) and genotype groups contained mixed males and females. Despite the range of ages utilized, every group was balanced to contain animals within the same age range to account for potential variability due to age and/or sex. Animal sample sizes are indicated in each figure. All data points, except for synapse density analysis, correspond to average values of multiple images analyzed for each mouse and represent single data points per mouse. All animal care and sacrifice procedures were approved by the University of Minnesota Institutional Animal Care and Use Committee (IACUC) in compliance with the National Institutes of Health guidelines for the care and use of laboratory animals under the approved animal protocol 2007-A38316.

### Immunohistochemistry

Sample preparation was performed as previously described [57]. Brains were cryo-sectioned into 16 μm coronal sections and stored in a 50% glycerol-50% tris-buffered saline (TBS) (25 mM Tris-base, 135 mM NaCl, 3 mM KCl, pH 7.6) solution at -20^○^C. For each experiment, three sections were used per mouse. Sections were blocked at room temperature in TBS with 0.2% Triton X-100 (TBST) for 1 hour, then incubated in primary antibodies overnight at 4C in TBST containing 5% NGS. Sections were blocked for 1 hour with secondary Alexa-fluorophore conjugated antibodies (Invitrogen) (1:200 in TBST with 5% NGS) at room temperature. Slides were mounted in Prolong Gold Antifade with DAPI (Invitrogen) and imaged. The following primary antibodies were used at the specified dilutions: GFAP (rabbit, PA1-10019, 1:1000), EM48 (mouse, mab3574, 1:200), and DARPP-32 (rat, mab4230,1:1000), VGLUT1 (guinea pig, ab5905, 1:1000), PSD-95 (rabbit, 51-6900, 1:500), NeuN (mouse, mab377, 1:1000), and Iba1 (rabbit, 011-27991, 1:500).

### DAB Immunostaining

Brains were cryo-sectioned into 16 μm coronal sections and stored in a 50% glycerol-50% tris-buffered saline (TBS) (25 mM Tris-base, 135 mM NaCl, 3 mM KCl, pH 7.6) solution at -20^○^C. For each experiment, three sections were used per mouse. Antigen retrieval was performed. Sections were blocked at room temperature in TBS with 0.3% Triton X-100 (TBST) containing 10% NGS for 1 hour, then incubated in primary antibodies overnight at 4C in TBST containing 5% NGS. Sections were incubated in secondary antibodies (Biotin) in TBST containing 5% NGS for 1 hour, then blocked in 3% hydrogen peroxide for 20 minutes. Sections were incubated in tertiary antibodies (VECTASTAIN Elite ABC HRP kit, Vector Laboratories) in TBST containing 5% NGS for 1 hour. Sections were incubated in DAB chromagen for 10 minutes. Sections were incubated in 0.1% Cresyl violet for 11 minutes. Slides were mounted in Prolong Gold Antifade with Permount (Fisher Scientific) and imaged. The following primary antibodies were used at the specified dilutions: EM48 (mouse, mab3574, 1:500), rbHtt (rabbit, ab109115,1:1000), 1C2 (mouse, mab5174, 1:500), and MAB2166 (mouse, mab2166, 1:500). 20x images of DAB stained CA1 hippocampal and cm dorsal striatal tissue were acquired on an epi-fluorescent microscope (Echo, Revolve). Blinded mages were opened in FIJI/ImageJ. For hippocampal sections, the Region of Interest (ROI) feature was used to outline the CA1 pyramidal cell layer. The Clear Outsides function was used to remove signal outside of the specified ROI. DAB and Cresyl violet staining were separated using the Colordeconvolution2 plugin, with settings normalized to zQ175 tissue. The DAB channel was thresholded (0-70) to include aggregates, which were counted using the Analyze Particles feature. Puncta counts were normalized per mm^2^. Mouse averages were calculated by averaging data obtained from images captured from three independent brain slices (n = 3 images/region/mouse).

### Imaging and cell counting analyses

For DARPP-32 quantification, 20x images were acquired on an epi-fluorescent microscope (Echo, Revolve). Using images obtained from representative zQ175 sections, background signal was determined by averaging fluorescence intensity from striatal white matter tracts and subsequently subtracted. After this subtraction, the average minimum and maximum fluorescence values of representative zQ175 sections were found using the Brightness/contrast feature and averaged. The brightness/contrast of each image was adjusted to 158 - 2609 (the average minimum and maximum values determined). 1 MSN with “borderline” DARPP-32 signal was outlined from each of 63 representative zQ175 images and its mean fluorescence signal intensity was measured. These 63 measurements were averaged to obtain the minimum mean fluorescence signal intensity necessary for an outlined cell to be considered DARPP-32^+^. WT, zQ175, and Q175DN images were blinded and opened in FIJI/ImageJ. All DARPP-32^+^ cells were manually outlined using the Region of Interest (ROI) feature and added to the ROI Manager. Images were adjusted using the methods specified above. The mean fluorescence signal intensity was measured for each ROI. All ROIs meeting or exceeding the threshold mean intensity value were counted as DARPP-32^+^. For each striatal section, 3 representative images were captured from the dorsal striatum and analyzed. Mouse averages for DARPP-32^+^ cell counts were determined by averaging estimates obtained from three sections (n = 9 images/mouse).

For GFAP analyses, 20x images of hippocampus were acquired on an epi-fluorescent microscope (Echo, Revolve) as described above. Blinded images were opened in FIJI/ImageJ. DAPI and GFAP channels were merged. Cells were manually counted using FIJI/ImageJ’s cell counter plugin. Cells were counted as GFAP^+^ only when a nucleus was detected (DAPI positive), as previously described [58]. GFAP^+^DAPI^+^ dual-positive cells on the perimeter of image were counted. For each section, three images were captured, corresponding to the CA1, CA2/3, and dentate gyrus regions, were captured. For each hippocampal section, 3 representative images were analyzed for each of the CA1, CA2/3, and dentate hippocampal regions. For each hippocampal region, mouse averages for GFAP^+^DAPI^+^ cell counts were determined by averaging estimates obtained from three sections (n = 9 images/region/mouse). For striatum analyses, 20x tiled GFAP and EM48 fluorescent images were acquired with a 16 μm z-dimension and 1 μm step size on a confocal microscope (Stellaris, Leica). Blinded images were opened in FIJI/ImageJ and condensed using Z project feature at maximum intensity, as previously described [58]. GFAP and EM48 images were merged. Cells were manually counted using FIJI/ImageJ’s cell counter plugin. Cells were counted as GFAP^+^ only when a nucleus was detected (DAPI positive). GFAP^+^DAPI^+^ dual-positive cells on the perimeter of image were counted. For each section, four images, corresponding to the cl, cm, dl, and dm striatal subregions, were captured using the same area per image (0.4 mm^2^). For each mouse, striatal subregion cells count was averaged from three sections (n = 3 images/subregion/mouse).

For Iba1^+^ cell counting analyses in the hippocampus and striatum, the cell counter plugin from ImageJ software was used and cells were counted manually. We used confocal images which were acquired with a 16 µm z-dimension and 1 µm step size, condensed using the Z Project function on Image J at max intensity. Cells or fascicles on the border of images were counted. All images used to calculate whole striatum cell counts were taken sampling throughout the striatum using the same area per image (0.4 mm^2^).

For DARPP-32 analyses, 20x images of the cm, cl, dm, and dl were acquired on an epi-fluorescent microscope (Echo, Revolve). Blinded images were opened in FIJI/ImageJ and thresholded to exclude white matter tracts. Mean fluorescence signal intensity (expressed in arbitrary units) within the thresholded pixels was recorded using FIJI/ImageJ’s measure function. For correlation analyses, EM48 puncta counts were normalized to the area of the thresholded pixels. For each brain section, one representative image was captured for each striatal region. Mouse averages were calculated by averaging data obtained from images captured from three independent brain slices (n = 3 images/region/mouse).

For HSF1 analyses, 20x images of hippocampus were acquired on an epi-fluorescent microscope (Echo, Revolve). Blinded images were opened in FIJI/ImageJ. CA1, CA3, and DG granule cell layers were manually outlined along DAPI-labeled cytoarchitecture and saved as ROIs. The hippocampal hilus was similarly outlined as the region between the DG granule cell layer. ROIs were opened on the corresponding HSF1 image. Mean fluorescence signal intensity (expressed in arbitrary units) within the ROI was recorded using FIJI/ImageJ’s measure function. For each brain section, one representative image was captured for each hippocampal region. Mouse averages were calculated by averaging data obtained from images captured from three independent brain slices (n = 3 images/region/mouse).

### HTT aggregate analyses

20x images of the hippocampus and dorsal striatum were acquired on an epi-fluorescent microscope (Echo, Revolve) as described above. Images were opened in FIJI/ImageJ. Background signal was subtracted and brightness/contrast was thresholded to 165 - 2149. Using the Region of Interest (ROI) feature, the granule cell layer (DG) or pyramidal cell layer (CA1, CA2/3) were outlined. The Clear Outsides function was used to remove signal outside of the specified ROI. Images were converted to RGB color and EM48 aggregates within the ROI were quantified blindly using the Puncta Analyzer plugin (Durham, NC, USA) on FIJI/ImageJ with size (pixels²) set to 3-500 and thresholded to the following: 70-255 (dorsal striatum), 65-255 (CA1), 85-255 (CA2/3), and 71-255 (DG). Dorsal striatum and CA1 threshold settings were set using images acquired from zQ175 tissue. CA2/3 and DG threshold settings were set using images acquired from Q175DN tissue, as zQ175 tissue contained insufficient aggregate load to accurately quantify EM48 puncta. To quantify cytoplasmic EM48 aggregation, images were thresholded to mask DAPI nuclei. EM48 quantification was performed as previously described to determine the number of cytoplasmic and nuclear aggregates. EM48 quantification analyses were conducted blindly with 3 separate sections analyzed per mouse. For each hippocampal section, 3 representative images were analyzed for each of the CA1, CA2/3, and DG hippocampal subregions. For each hippocampal subregion, mouse averages for number and size (pixels²) of EM48 aggregates were determined by averaging estimates obtained from three sections (n = 9 images/region/mouse). For each striatal section, 3 representative images were analyzed for the dorsal striatum and mouse averages for the number of EM48 aggregates were determined by averaging estimates obtained from the three sections (n = 9 images/mouse).

To analyze EM48 aggregation in the dm, dl, cm, and cl striatal subregions, 20x tiled EM48 fluorescent images from the striatum were acquired with a 16 μm z-dimension and 1 μm step size on a confocal microscope (Stellaris, Leica). Images were blinded, opened in FIJI/ImageJ, and condensed using the Z project feature at maximum intensity. Images were converted to RGB color and EM48 aggregates were quantified using the Puncta Analyzer plugin (Durham, NC, USA) on FIJI/ImageJ with threshold set to 77-255 and size (pixels²) set to 3-500. Threshold settings were set using images acquired from ZQ175 dl striatal tissue. Cytoplasmic EM48 aggregation was quantified using thresholded images to mask DAPI nuclei. EM48 quantification analyses were conducted blindly. Images were analyzed in their entirety. Per striatal section, 1 representative image was analyzed for each of the dm, dl, cm, and cl striatal subregions; thus, for each striatal subregion, mouse averages for number and size (pixels²) of EM48 aggregates were determined by averaging estimates obtained from 3 images.

### VGLUT1/PSD-95 colocalization quantification of synaptic density

Images for synaptic density analysis were acquired on a Leica Stellaris 8 confocal microscope (optical section depth 0.34 mm, 15 sections per scan, bit depth 12) in the molecular cell layer of CA1 and dentate gyrus in the hippocampus, and in the synaptic zone in the dorsal striatum at 63X magnification. Each image was processed using Leica Lightning Processing in the LAS-X software prior to export. Maximum projections of 3 consecutive optical sections were generated. Significant group differences were calculated using *n* = 3 consecutive optical sections per animal, n = 3 measurements per coronal section, and a total of n = 3 animals per genotype resulting in a total of 18 data points per genotype. Data was averaged per coronal section resulting in 3 data points per mouse and 9 data points per genotype. Puncta analyses were conducted blinded using the PunctaAnalyzer Plugin (Durham, NC, USA) on ImageJ, as previously described [57] following recommendations by Ravaliat et al. [59].

### RNA preparation and RT-qPCR

RNA was extracted from microdissected hippocampal tissues using the RNeasy extraction kit (Qiagen; Germantown, MD, USA) according to the manufacturer’s instructions. cDNA was prepared using the Superscript First Strand Synthesis System (Invitrogen; Waltham, MA, USA) according to the manufacturer’s instructions. SYBR green-based qPCR was performed with SYBR mix (Genesee; EL Cajon, CA, USA) using the LightCycler 480 System (Roche; Basel, Switzerland). Primers used are as follows: GAPDH (Forward: 5’-ACACATTGGGGGTAGGAACA-3’, Reverse: 5’-AACTTTGGCATTGTGGAAGG-3’), Hsp25 (Forward: 5’-CGAAGAAAGGCAGGATGAAC-3’, Reverse: 5’-CGCTGATTGTGTGACTGCTT-3’), Hspa1a (Forward: 5’-TGGTGCTGACGAAGATGAAG-3’, Reverse: 5’-AGGTCGAAGATGAGCACGTT-3’). Htt (Forward: 5’-TGCAGCCTCTGTGAAGAGTG-3’, Reverse: 5’-TGGGATCTAGGCTGCTCAGT -3’). Each sample was tested in triplicate and normalized to GAPDH levels. For analysis, the 2^-ΔΔCt^ method was used to calculate the relative for gene expression.

### NeuN Hippocampal Layer Thickness

20X hippocampal tiles were acquired using a confocal microscope with a 16 μm z-dimension and 1 μm step size (Leica Stellaris 8). Images were opened in Imaris image analysis software (Oxford Instruments). A batch process was initiated allowing for the application of set parameters to all images. Using the ellipse tool an area around CA1 and CA3 as defined by Allen Brain Atlas images (adult mouse coronal, P56 and P55). Area was applied to all images and standardized by pixel number. Three lines were randomly drawn across the width of the area previously assigned to CA1 and CA3 and the angles were adjusted to match the angle of dendrites in the molecular layer of the hippocampus. The widths of these lines were averaged in order to attain the average width per image for CA1 and CA3.

### Quantification and statistical analysis

Data from 3 brain sections per animal were averaged to generate an animal average, subsequently used in statistical analysis, except in synaptic density analyses where average per coronal section was used. Sample sizes for each experiment are specified in the relevant figure legend. Data are expressed as Mean ± SEM, analyzed for statistical significance, and displayed by Prism 9 software (GraphPad, San Diego, CA, USA). The accepted level of significance was p ≤ 0.05. All p-values are shown in Table 1.

**Table 1.**
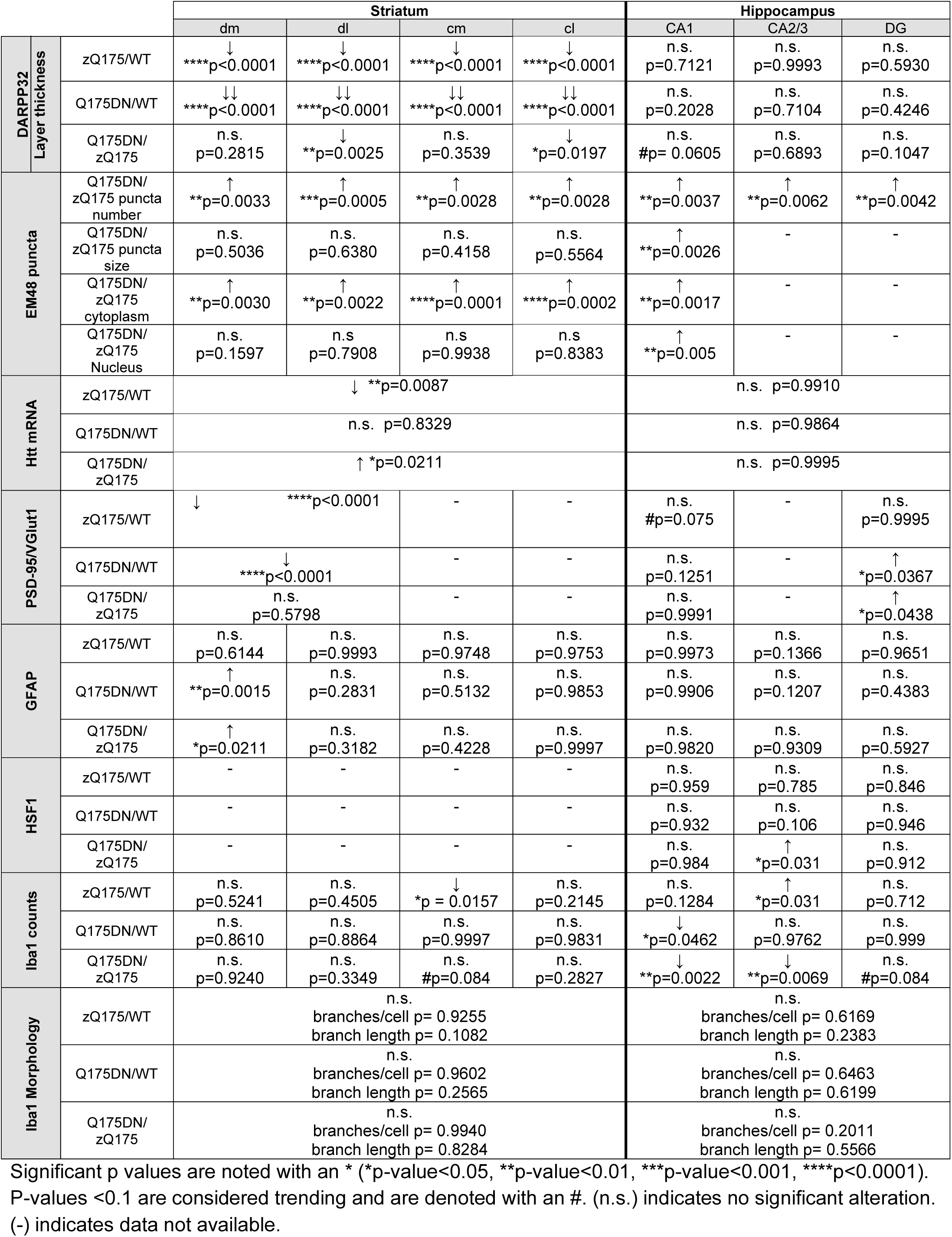
Summary of results and p values.

## RESULTS

### Excision of the PGK-Neomycin cassette in zQ175 (C57BL/6 background) results in increased mHTT aggregation in the striatum and hippocampus

Enhanced neuropathology has been observed in specific regions of the striatum in various HD mouse models, especially in the dorsal areas [58]. Therefore, we conducted HTT aggregation analyses in four subdivisions of the striatum corresponding to the dorsomedial (dm), dorsolateral (dl), centromedial (cm) and centrolateral (cl) (**Figure 1A**) to see if previously reported regional selectivity in striatal pathology was associated with enhanced HTT aggregation. We evaluated HTT aggregation in zQ175 and Q175DN mice via immunohistochemistry and DAB staining using different HTT antibodies (**Supplementary Figure 2, 3**) and selected the use of EM48 as the preferred antibody to assess HTT aggregation [42, 58, 60]. This antibody recognizes the N-terminal (aa1-212) of HTT exon1 [61] and has been extensively used to detect mHTT aggregates by immunohistochemistry [41, 62–64]. Previous reports have indicated increased HTT aggregation in the striatum of Q175FDN mice compared to control Q175F [45]. Since striatal degeneration in HD has been shown to be more prominent in dorsal regions of the striatum, we decided to assess whether HTT aggregation varied among regions of the striatum in either genotype. We found that Q175DN presented enhanced levels of HTT aggregation compared to zQ175 in all striatal subregions. EM48 DAB staining showed highly comparable results with EM48 immunofluorescence (**Supplementary Figure 2A, B**). HTT aggregation was increased across the cl (p = 0.0033), cm (p = 0.0005), dl (p = 0.0028), and dm (p = 0.0028) striatum although no differences among regions were observed in either zQ175 or Q175DN mice (**Figure 1B, C**). Despite the enhancement in the number of EM48^+^ puncta in Q175DN mice, no significant differences were observed in EM48^+^ puncta size in any striatal subregion compared to zQ175 (**Figure 1D**). When assessing the distribution of EM48 aggregates between the cytoplasm and the nucleus of cells for each of the striatum subregions, only a significant increase was observed in the cytoplasmic content of EM48+ puncta (**Figure 1E**) but not in the nuclear where similar levels were observed between zQ175 and Q175DN (**Figure 1E**).

**Figure 1.**
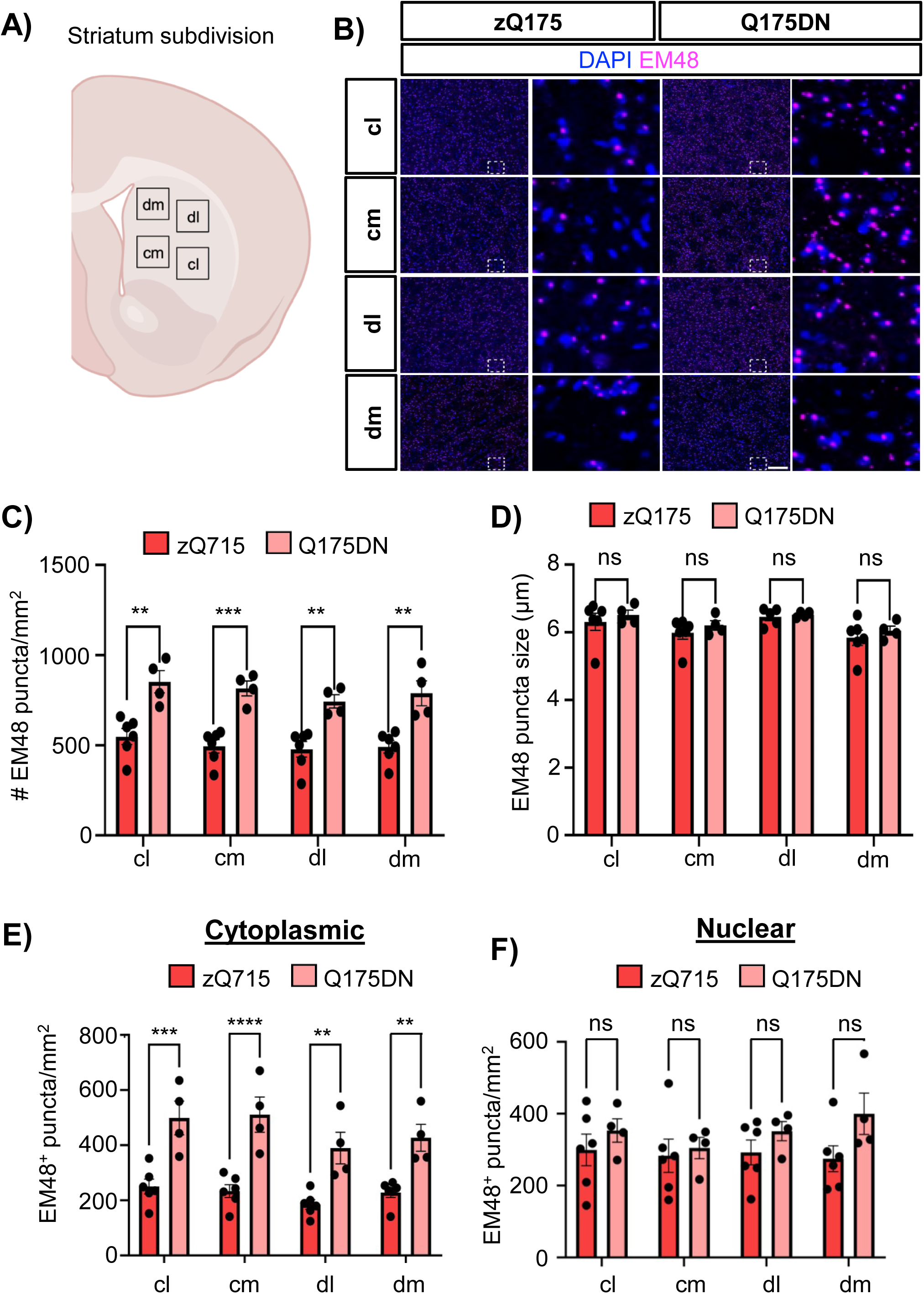
Q175DN presents enhanced mHTT aggregation across striatal subregions. (**A**) Graphical representation of image locations used in mHTT analyses in the striatum. (**B**) Representative DAPI and EM48 immunostainings in the cl (centrolateral), cm (centromedial), dl (dorsolateral), and dm (dorsomedial) striatum in zQ175 and Q175DN. Scale bar, 90 µm. (**C, D**) mHTT aggregation quantified by number of EM48^+^ puncta per mm^2^ (**C**) and EM48 puncta size in µm (**D**) in zQ175 and Q175DN in the cl, cm, dl and dm striatum (n=6 zQ175, n=4 Q175DN). One zQ175 outlier removed from dl EM48 puncta size quantification. (E, F) EM48^+^ puncta per mm^2^ in the cytoplasm (E) and nucleus (F). Error bars denote mean ± SEM. Two-way ANOVA with Sidak’s multiple comparisons in C and D. [(C) Interaction: F = 0.125, df = 3; striatal subregion: F = 1.290, df = 3; genotype: F = 81.31, df = 1. (D) Interaction: F = 0.06900, df = 3; striatal subregion: F = 3.612, df = 3; genotype: F = 1.653, df = 1; (E) Interaction: F = 0.4908, df = 3; striatal subregion: F = 2.417, df = 3; genotype: F = 75.98, df = 1, (F) Interaction: F = 0.5370, df = 3; striatal subregion: F = 0.3927, df = 3; genotype: F = 4.2828, df = 1]. \*\**p* < 0.01, \*\*\**p* < 0.001.

We also evaluated EM48 puncta (number and size) in three subdivisions of the hippocampus corresponding to the cornu ammonis (CA1 and CA2/3) and the dentate gyrus (DG) in zQ175 and Q175DN mice (**Figure 2A, B**). We observed a significant increase in the number of HTT aggregates in the molecular cell layer of CA1 (p = 0.0037) (**Figure 2C**), as well as a significant increase in number of mHTT aggregates in the molecular cell layer of CA2/3 (p = 0.0062) (**Figure 2D**) and the DG (p = 0.0042) (**Figure 2E**). Interestingly, when comparing the number of EM48+ aggregates between different hippocampal subregions for each genotype it was apparent that the highest increase in EM48+ aggregates for Q175DN occurred in the DG (**Supplementary Figure 4**). We also conducted analyses to assess if there was a differential distribution of aggregates between the cytoplasm and the nucleus of hippocampal cells. Since only the CA1 region showed a significant proportion of aggregates to be measured in the zQ175 we focused on further analyses in this region. We found that, contrary to the striatum, both the cytoplasmic and nuclear aggregates were significantly increased in the CA1 of Q175DN mice (**Figure 2F, G**). Interestingly, the number of aggregates in the cytoplasm were considerably low compared to those observed in the nucleus indicating that most aggregates in the CA1 correspond to nuclear aggregates revealing a nearly identical pattern as that observed in **Fig. 2C** for total EM48+ puncta. We also evaluated the size of the EM48^+^ puncta in the CA1 and observed a significant increase in the puncta size in Q175DN compared to zQ175 (**Figure 2F**). These findings suggest that the PGK-Neomycin cassette negatively affects *Htt* expression and that its excision leverages HTT expression and aggregation not only in the striatum but also in other brain regions such as the hippocampus where very little EM48 puncta is observed when the Neomycin cassette is present. When we assessed the levels of total *Htt* transcripts in the striatum of WT, zQ175, and Q175DN mice we observed a significant decrease in zQ175 compared to WT, as previously reported [65], and a significant increase in Q175DN compared to zQ175 (**Supplementary Figure 5A**). The enhanced accumulation of total *Htt* in Q175DN could be attributed to an enhanced expression of the transgenic m*Htt*. Interestingly, when the levels of total *Htt* transcripts were assessed in the hippocampus, no significant differences were observed among the different genotypes (**Supplementary Figure 5B**). Similar results have been obtained when assessing full length *Htt* transcripts in the hippocampus of WT and zQ175 mice using QuantiGene 10-multiplex and qPCR assays [65]. However, more complex transcript analyses using a custom-made 14-plex QuantiGene assay have captured differences in full length *Htt* in the hippocampus of zQ175. Nevertheless, our comparative analyses between striatum and hippocampus using qPCR are sufficient to reveal that *Htt* expression changes differ between these two brain regions and mouse models and imply that the enhanced aggregation of HTT observed in the hippocampus of Q175DN mice cannot be solely attributed to enhanced expression of *Htt* (endogenous or mutant).

**Figure 2.**
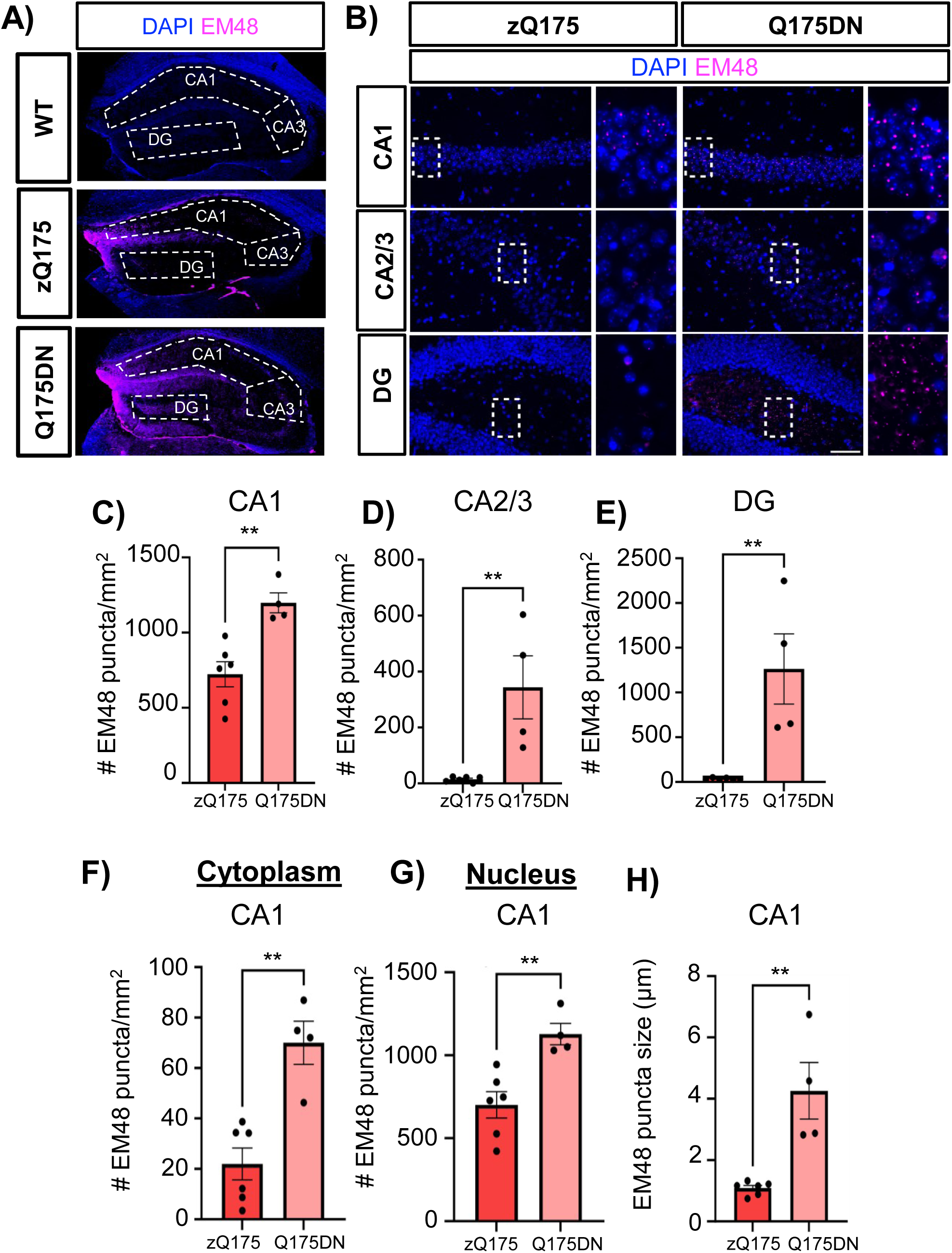
Q175DN presents enhanced regional differences in mHTT aggregation in the hippocampus. (**A**) Representative DAPI and EM48 coronal immunostainings of the whole hippocampus in WT, zQ175, and Q175DN. Hippocampal regional image locations used in mHTT analyses are indicated. (**B**) Representative DAPI and EM48 of the CA1, CA2/3 and DG. Scale bar, 50 µm. The white dashed rectangles delineate a representative region of each hippocampal subregion shown next to the image with a higher magnification. (**C-E**) Hippocampal mHTT aggregation quantified by number of EM48^+^ puncta per mm^2^ in Q175DN mice compared to zQ175 mice in the CA1 (**C**), CA2/3 (**D**) and DG (**E**) (n=6 zQ175, n=4 Q175DN). (**F, G**) EM48^+^ puncta per mm^2^ in Q175DN mice compared to zQ175 mice in the cytoplasm (**F**) and nucleus (**G**) of CA1 hippocampal cells. (**H**) EM48 puncta size in µm in the CA1 of Q175DN and zQ175 mice. Error bars denote mean ± SEM. Un-paired Student’s *t*-test in C-F. [(C) *t* = 3.954, df = 8; (D) *t* = 3.720, df = 8; (E) *t* = 5.419, df = 8; (F) *t* = 4.633, df = 8; (G) *t* = 3.829, df = 8; (H) *t* = 4.293, df = 8]. \*\**p* < 0.01, \*\*\**p* < 0.001.

### Enhanced mHTT aggregation in Q175DN correlates with enhanced neuronal dysregulation in the striatum but not in the hippocampus

Dopamine- and cAMP-regulated phosphoprotein 32 (DARPP-32) is an important dopamine signaling protein that is critically involved in regulating electrophysiological, transcriptional, and behavioral responses. DARPP-32 is highly abundant in MSNs and its expression progressively declines in HD [41, 66, 67]. Decreased expression of DARPP-32 in HD MSNs is considered a marker of transcriptional dysregulation and neuronal degeneration [68]. We first evaluated the immunoreactivity of DARPP-32 in WT, zQ175 and Q175DN in the whole striatum and the number of DARPP-32^+^ cells was quantified. A significant and overt decrease in DARPP-32 was observed in zQ175 when compared to WT, as previously reported [41, 57] (**Figure 3A, B**). More importantly, we found that the number of DARPP-32^+^ cells in the Q175DN significantly decreased compared to zQ175 mice indicating a worsening in striatal degeneration (**Figure 3A, B**). The number of DARPP-32^+^ cells was inversely correlated to the number of EM48^+^ puncta found in the striatum (**Figure 3C**). These results were in line with previous observations in the Q175FDN mouse model [45]. However, despite differences in mHTT aggregation and neuronal degeneration between zQ175 and Q175DN mice, no changes in body weight were observed (**Supplementary Figure 6**).

**Figure 3.**
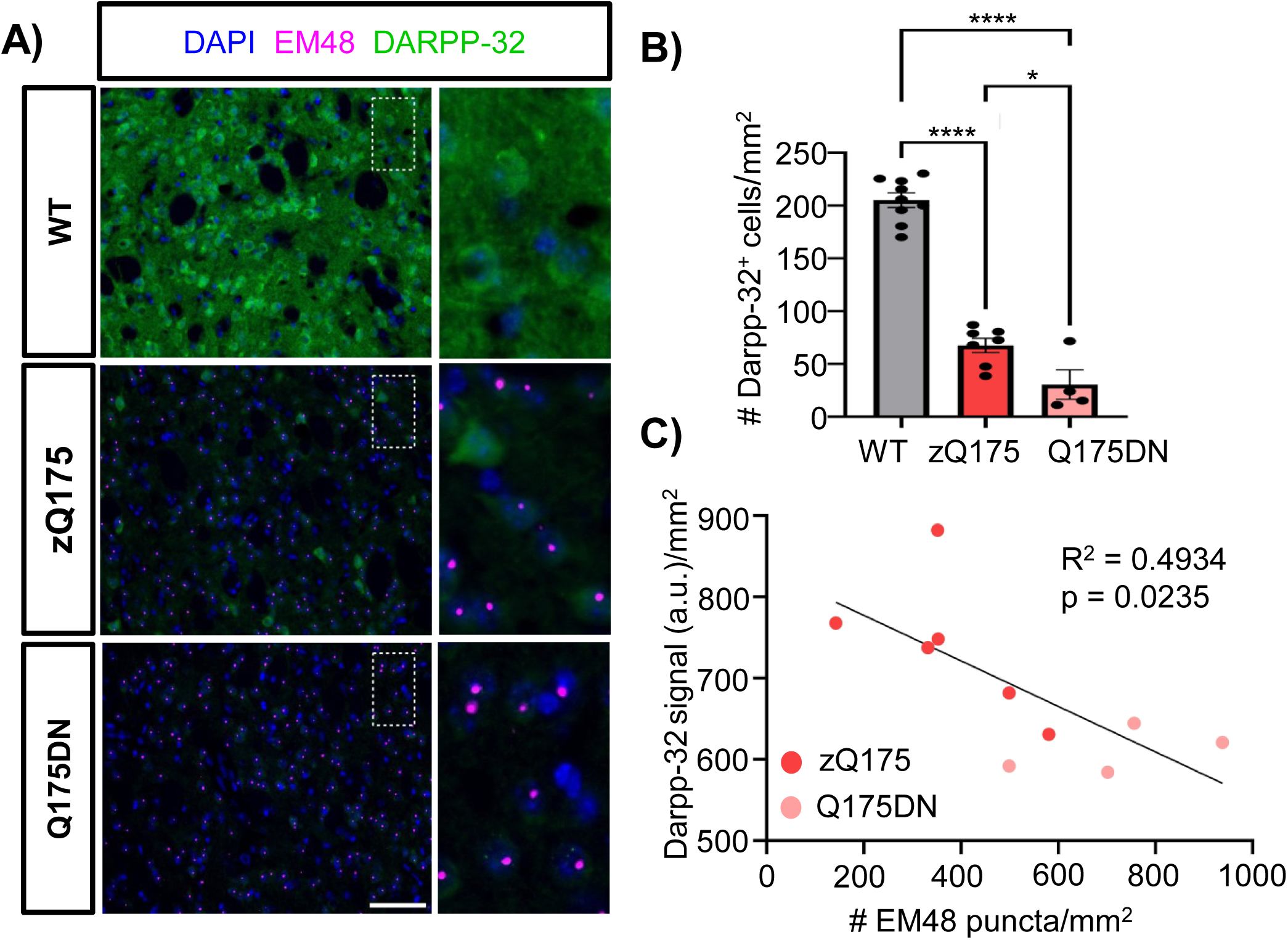
Enhanced susceptibility to striatal neuron degeneration in Q175DN mice. (**A**) Representative immunostaining of DARPP-32 and EM48 in the dorsal striatum of WT, zQ175, and Q175DN mice. Scale bar, 50 µm. (**B**) Quantification of the number of DARPP-32^+^ cells per mm^2^ in WT (n=9), zQ175 (n=7), and Q175DN (n=4). Error bars denote mean ± SEM. One-way ANOVA with Tukey’s post hoc test. (**C**) Correlation analysis of DARPP-32^+^ signal (a.u)/mm^2^ versus number of EM48^+^ puncta/mm^2^. Data from zQ175 (n=7) and Q175DN (n=4) mice. R^2^ represents the coefficient of determination. Simple linear regression. [(B) F = 128.8, df = 2; (C) F = 7.791, df = 10]. \**p* < 0.05, \*\*\*\**p* < 0.0001.

We also evaluated the levels of DARPP-32 in each of the striatum regions (dm, cm, dl, and cl) to determine if specific subregional degeneration was observed, and we performed correlation analyses between DARPP-32 and EM48 for each region (**Supplementary Figure 7**). We found that DARPP-32 signal was lower in the dm and cm compared to the dl and cl in the zQ175 mice (**Supplementary Figure 7A**). This suggested a greater impact in medial areas of the striatum compared to lateral areas. When looking at the Q175DN mouse the levels of DARPP-32 were reduced across all areas of the striatum with no significant differences between any of the striatal regions, suggesting an enhanced degeneration that expanded to more lateral areas of the striatum. Correlation analyses also revealed a significant inverse correlation between DARPP-32 signal and EM48 levels in the dm (p=0.049) and cm (p=0.001) and trends in the dl (p=0.099) and cl (p=0.058) (**Supplementary Figure 7B-E**).

We then evaluated whether the enhanced HTT aggregation observed in the hippocampus of Q175DN also increased neuronal degeneration in this brain region. For this, we used the neuronal marker NeuN and we evaluated the thickness of the granule cell layer in the CA1, CA3 and DG. First, we did not observe any signs of hippocampal layer atrophy between WT and zQ175 mice in any of the tested hippocampal regions (**Figure 4A-D**). Surprisingly, no significant differences in granule cell layer thickness between zQ175 and Q175DN were observed in any hippocampal regions (**Figure 4B-D, Supplementary Figure 8**) despite the significant difference observed in EM48^+^ aggregates between these two genotypes (**Figure 2C-E**). These findings suggested an enhanced hippocampal spare capacity to the toxicity of mHTT that contrasts with the enhanced neuronal dysregulation experienced by striatal medium spiny neurons.

**Figure 4.**
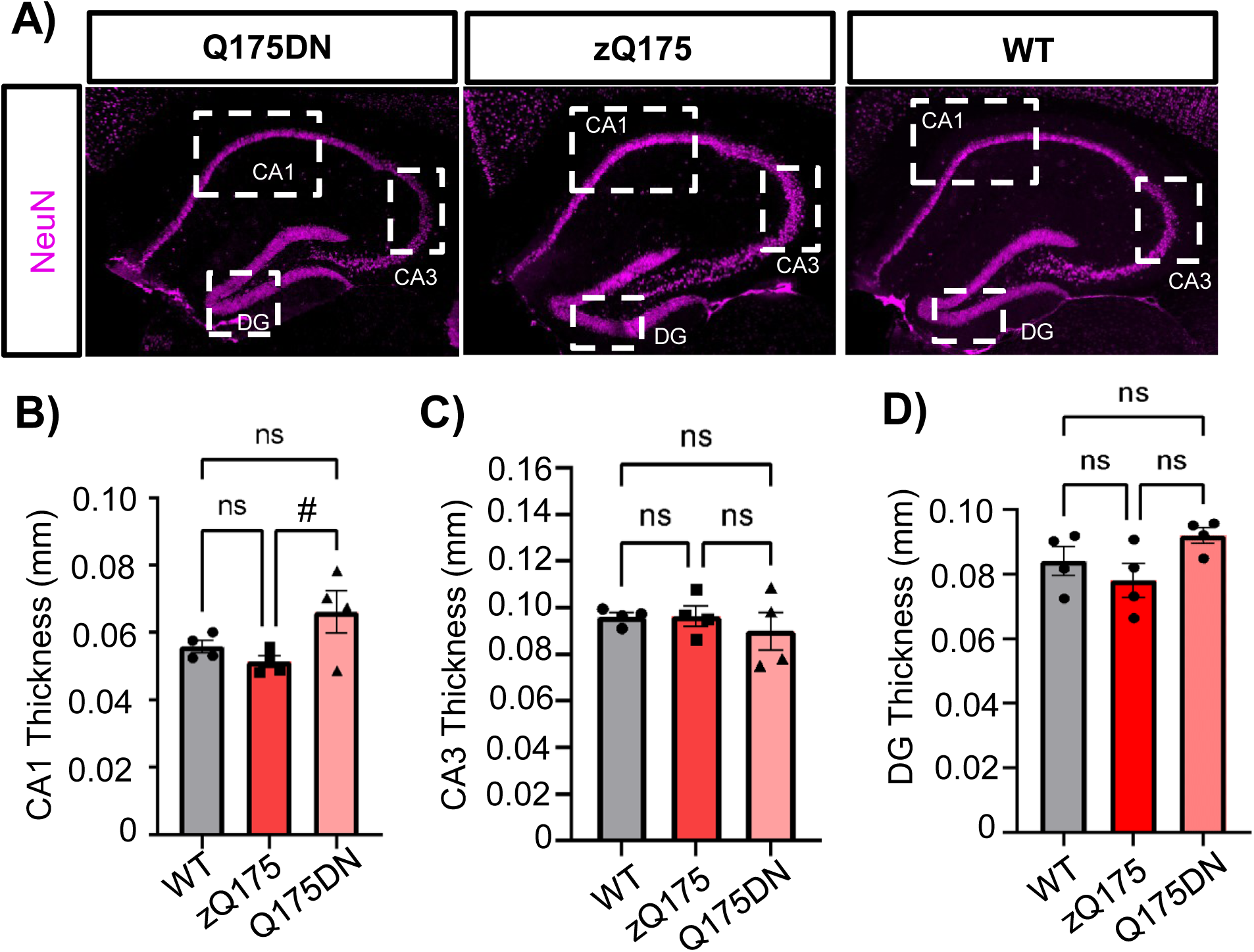
Hippocampal layer thickness is not affected in Q175DN mice. (**A**) Representative immunostaining of NeuN in the hippocampus of WT, zQ175, and Q175DN mice. Scale bar, 100 µm. (**B-D**) Granule cell layer thickness (μm) in the CA1 (**B**), CA3 (**C**) and DG (**D**) of WT (n=4), zQ175 (n=4), and Q175DN (n=4) mice. Error bars denote mean ± SEM. One-way ANOVA with Tukey’s post hoc test. #*p* < 0.1. [(B) F = 3.754, df = 11; (C) F = 6.075, df = 11; (D) F = 2.694, df = 11].

### Glial responses are differentially altered in the Q175DN hippocampus

Gliosis is a reactive process occurring after most types of central nervous system injuries and neurodegeneration and is the result of focal proliferation and phenotypic alteration of glial cells. Neuronal dysfunction and degeneration in HD are accompanied by increased astrocyte density and astrocyte pathology in both mouse models and patients [9, 69, 70]. The expression of the glial fibrillary acidic protein (GFAP) is considered as marker of reactive astrocytes [71], although it is also naturally expressed in high levels in astrocytes associated with white matter [72]. We recently showed an age-dependent increase in the number of GFAP^+^ cells in the striatum of zQ175 mice compared to WT, especially within the dm striatum, that becomes more apparent in old animals (∼18 months) [58]. We evaluated whether enhanced mHTT aggregation in Q175DN mice would enhance the accumulation of GFAP^+^ cells at younger ages in the dm or in other striatal regions. We therefore quantified the number of GFAP^+^ cells in the cl, cm, dl, and dm striatum between WT, zQ175 and Q175DN mice (**Figure 5A, B**). As previously reported [58], the number of GFAP^+^ cells in both WT and zQ175 were very low in central regions of the striatum (cl and cm) with higher number in the dorsal striatum (dl and dm) (**Figure 5B**). Although no significant differences were observed in the number of GFAP^+^ cells in any of the tested regions between WT and zQ175 at 12 months of age, we observed a significant increase in the dm striatum of Q175DN mice compared to WT and zQ175 (**Figure 5 B**). The enhanced astrogliosis in the dm of Q175DN coincides with a greater correlation between decreased DARPP-32 levels and increased mHTT aggregation compared to zQ175 mice.

**Figure 5.**
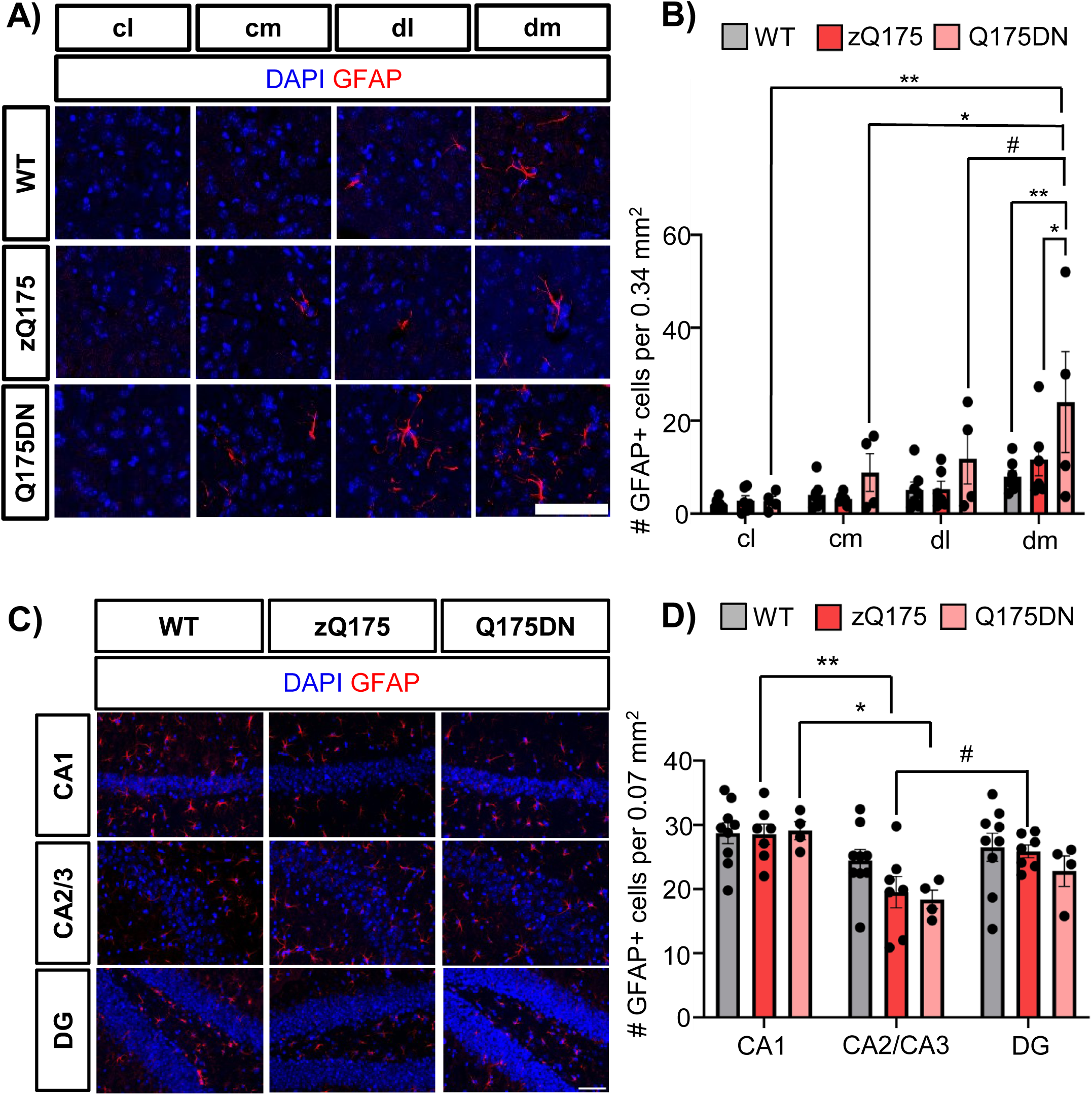
Number of GFAP^+^ astrocytes increased in the dorsal striatum of Q175DN mice but not in the hippocampus. (**A, B**) Representative immunostainings of GFAP in the cl (centrolateral), cm (centromedial), dl (dorsolateral), and dm (dorsomedial) striatum of WT, zQ175, and Q175DN mice. DAPI stains nuclei. Scale bar, 50 µm. (**B**) Quantification of GFAP^+^ cells per 0.34 mm^2^ in the cl, cm, dl and dm striatum of WT (n=7), zQ175 (n=6), and Q175DN (n=4). (**C**) Representative immunostainings of GFAP and DAPI in the hippocampus of WT, zQ175, and Q175DN mice. Scale bar, 50 µm. (**D**) Quantification of GFAP^+^ cells per mm^2^ in the CA1, CA2/3, and DG hippocampal regions of WT (n=9), zQ175 (n=7), and Q175DN (n=4). Error bars represent mean ± SEM. Two-way ANOVA with Sidak’s multiple comparisons. [(B) Interaction: F = 1.192, df = 6; striatal subregion: F = 8.781, df = 3; genotype: F = 5.679, df = 2. (D) Interaction: F = 0.9015, df = 4; hippocampal subregion: F = 11.37, df = 2; genotype: F = 1.858, df = 2]. #*p* < 0.1, \**p* < 0.05, \*\**p* < 0.01.

We next evaluated astrocyte pathology in the hippocampus (**Figure 5C, D**). First, we found that the abundance of GFAP^+^ astrocytes in all regions of the hippocampus were higher than in the striatum for all genotypes (**Supplementary Figure 9**) and presented a more homogeneous distribution. No differences were found in the amount of GFAP^+^ cells between the different hippocampal regions for WT mice, although zQ175 and Q175DN presented lower GFAP^+^ cell in the CA2/3 region compared to the CA1 and DG. Despite seeing enhanced HTT aggregation in the Q175DN in all areas of the hippocampus, we observed no significant differences in GFAP^+^ cell number compared to zQ175 in any of the tested regions (**Figure 3C, D**).

Increased reactive microglia has also been reported in the striatum, cortex, and globus pallidus of HD patients in all grades of pathology and their number correlates with the degree of neuronal loss in the striatum [73, 74]. However, the recapitulation of microgliosis across different HD mouse models has been inconsistent [42, 75–78]. We used the ionized calcium binding adaptor molecule 1 (Iba1) as a marker for detecting reactive microglia [79] and quantified the number of Iba1^+^ cells throughout the striatum and hippocampus (**Figure 6A, B**). Except for the cm striatum, where surprisingly we found significantly fewer Iba1^+^ microglia in zQ175 mice compared to WT mice, no other significant differences in Iba1^+^ cell counts were found between genotypes (**Figure 6B**). This data is consistent with previous reports showing the lack of significant microgliosis between WT and symptomatic zQ175 in the dorsal striatum [42] and suggests that the number of Iba1^+^ cells in the striatum does not correlate with mHTT aggregate load. Regarding the hippocampus, the average number of Iba1^+^ cells in zQ175 was higher than WT in all tested regions although it was only statistically significant in the CA2/3 region (**Figure 6C, D**). Contrary to what it was expected, Q175DN mice displayed a significant decrease in Iba1^+^ cell counts compared to zQ175 in the CA1 and CA2/3 and a trend towards decreased counts in the DG. When we assessed the morphology of Iba1+ microglia in the striatum (**Figure 7A-C**) and hippocampus (**Figure 7C-E**) we found no significant differences in either the number of branches or branch length for any of the genotypes. However, a trend towards decreased branch length was found for both zQ175 and Q175DN in the striatum, while a trend towards increased branch length was found in the hippocampus for zQ175 being lower in the Q175DN. Overall, this data suggests that microgliosis in the hippocampus might not necessarily respond to HTT aggregation.

**Figure 6.**
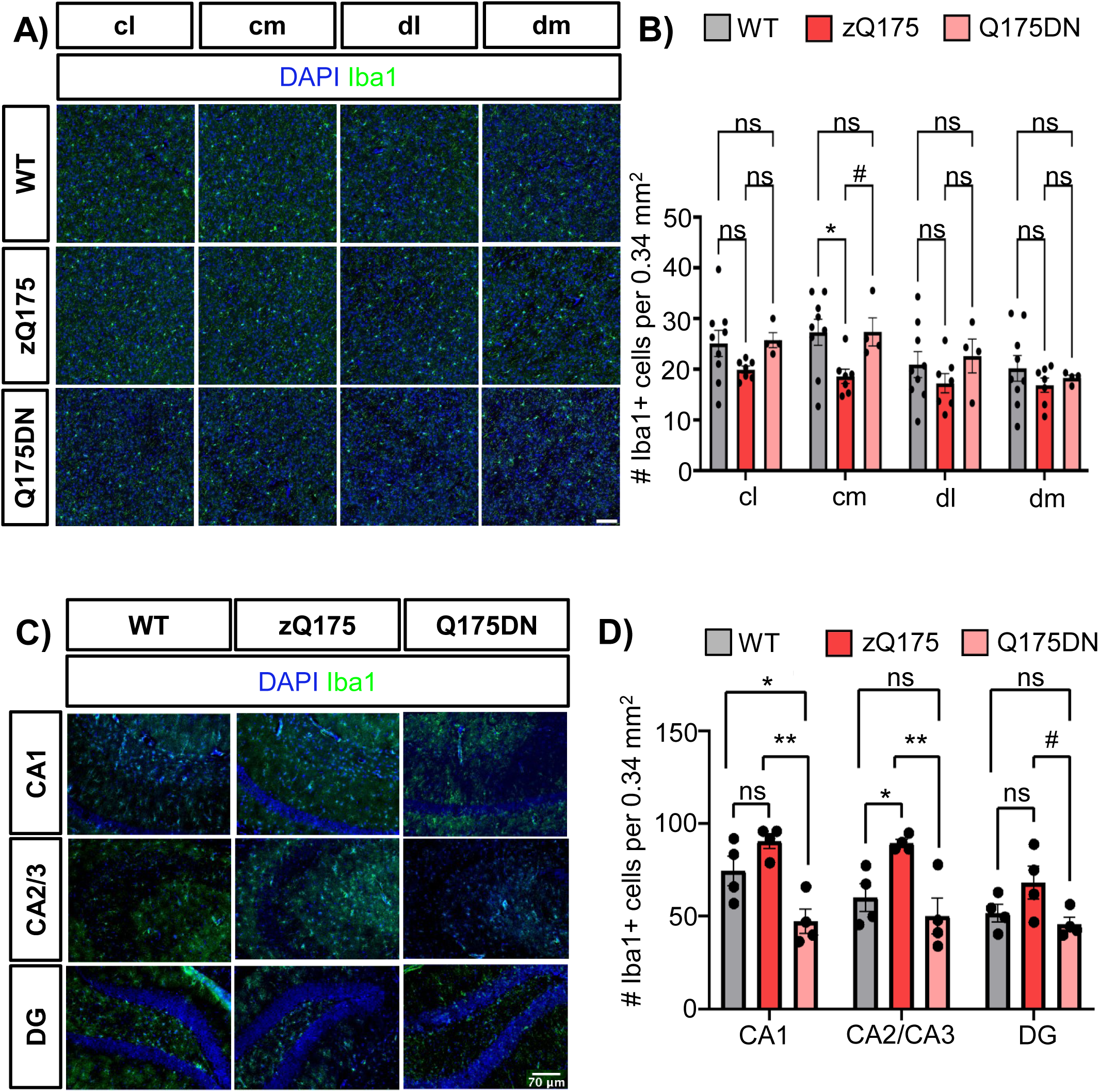
Microglia dysregulation in the Q175DN mouse model is increased in the dorsal striatum but reduced in the hippocampus. (**A**) Representative immunostainings of Iba1 in the cl (centrolateral), cm (centromedial), dl (dorsolateral), and dm (dorsomedial) striatum of WT, zQ175, and Q175DN mice. DAPI stains nuclei. Scale bar, 90 µm. (**B**) Quantification of Iba1^+^ cells per 0.34 mm^2^ from A in the cl, cm, dl, and dm striatum of WT (n=9), zQ175 (n=7), and Q175DN (n=4). (**C**) Representative immunostainings of Iba1 in the hippocampus of WT, zQ175, and Q175DN mice. DAPI stains nuclei. Scale bar, 70 µm. (**D**) Quantification of Iba1^+^ cells per 0.34 mm^2^ from C for WT (n=4), zQ175 (n=4), and Q175DN (n=4). Error bars represent mean ± SEM. One-way ANOVA with Tukey’s post hoc test. Two-way ANOVA with Sidak’s multiple comparisons. [(B) Interaction: F = 1.192, df = 6; striatal subregion: F = 8.781, df = 3; genotype: F = 5.679, df = 2. (D) Interaction: F = 0.4870, df = 6; hippocampal: F = 3.874, df = 3; genotype: F = 6.887, df = 2]. #*p* < 0.1, \**p* < 0.05, \*\**p* < 0.01.

**Figure 7.**
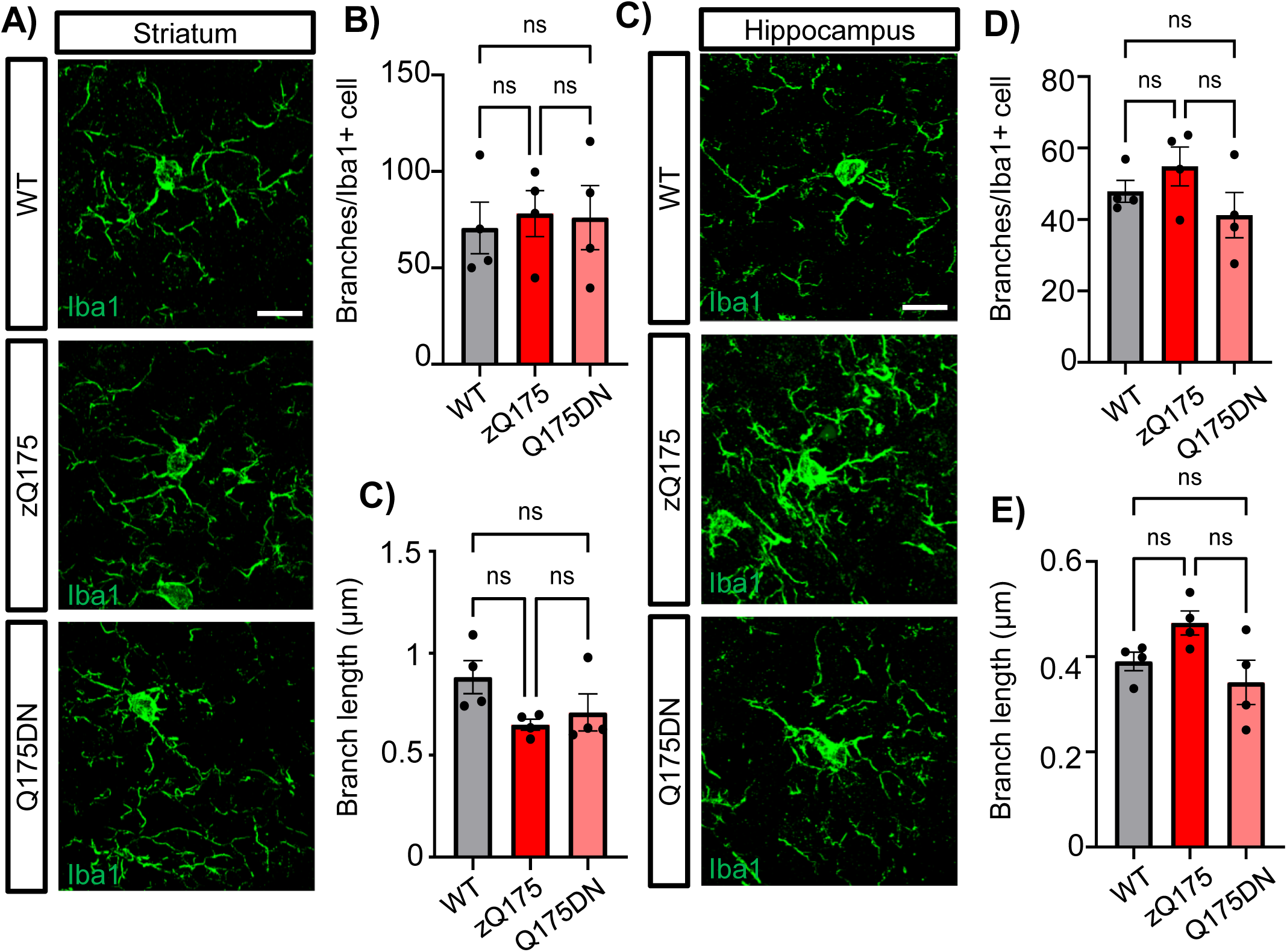
Morphology of microglia is unchanged between Q175DN and zQ175 mice in the hippocampus and striatum. (A) Representative immunostainings of Iba1 in the hippocampus of WT (n=4), zQ175 (n=4), and Q175DN (n=4). Scale bar, 5 µm. (B) Quantification of branches per Iba1+/cell in each 63x image. (C) Quantification of mean branch length in Iba1+ cells in µm. (D) Representative immunostainings of Iba1 in the striatum of WT (n=4), zQ175 (n=4), and Q175DN (n=4). Scale bar, 5 µm. (E) Quantification of branches per Iba1+/cell in each 63x image. (F) Quantification of mean branch length in Iba1+ cells in µm. Error bars denote mean ± SEM. One-way ANOVA with Tukey’s post hoc test in B,C and E,F. [(B) Interaction: F = 1.759, df = 11, (C) Interaction: F = 3.786, df = 11, (E) Interaction: F = .07551, df = 11, (F) Interaction: F= .664, df = 11.

### Synaptic density is increased in the hippocampus of Q175DN mice

We and others have previously shown that deficits in synaptic density are a robust parameter indicative of neuronal dysfunction and degeneration in the striatum [57, 80–83]. Striatal synaptic density declines during aging and such phenomenon is exacerbated in HD models [57, 80–83]. Synapses from both thalamo-striatal and cortico-striatal inputs are significantly decreased in zQ175 mice at symptomatic stages [57, 80, 84–87]. On the other hand, PET analyses have shown hippocampal synaptic alterations in both the zQ175 [43, 52, 88] and in Q175FDN and Q175DN mice [46, 47] although specific synapse density analyses to confirm such alterations are lacking. Based on the scarce information regarding hippocampal synaptic integrity in either zQ175 or Q175DN mice we investigated whether increased aggregation in the Q175DN corresponded to altered synaptic density in the hippocampus as compared to zQ175 mice. We measured synapse density by assessing the co-localization between the vesicular transporter VGlut1 (marker for excitatory cortical projections) and the postsynaptic protein PSD-95, as we previously reported [80] (**Figure 8**). We and others have reported a decrease in cortico-striatal synapse density in zQ175, especially at symptomatic stages [80, 84] and decided to explore if cortico-striatal synapse loss was aggravated by the enhanced accumulation of mHTT in Q175DN mice. Although we found a significant depletion in both PSD-95 and VGlut1 expression and colocalization in zQ175 compared to WT (**Figure 8B-D**), as we previously reported [80], no significant differences were observed between zQ175 and Q175DN mice.

**Figure 8.**
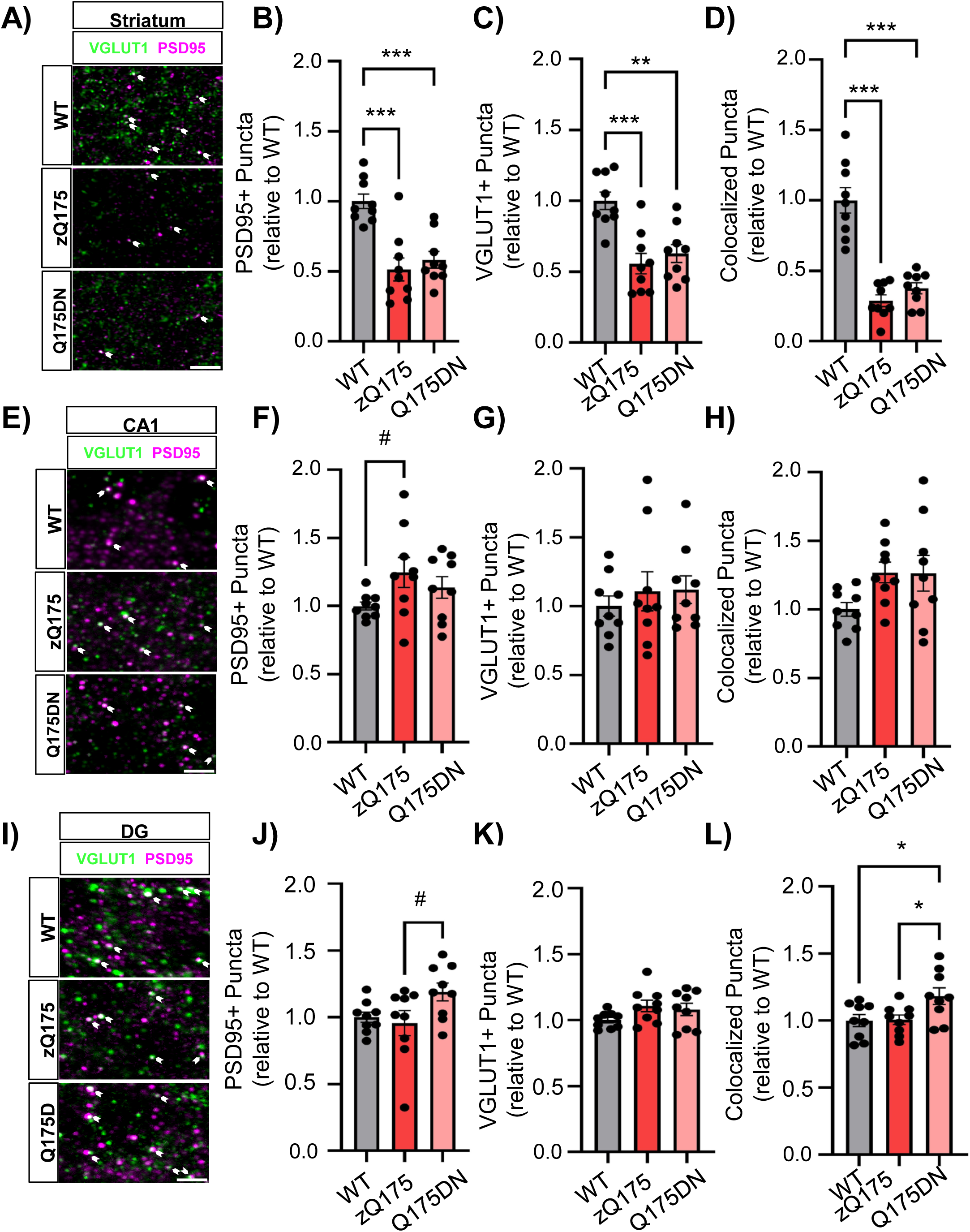
Synaptic density is increased in the DG region of the hippocampus of Q175DN mice. (**A**) Representative images from molecular cell layer of CA1. (**B**) Graphical representation of image location used in synaptic density analysis. (**C**) Synaptic density quantified by colocalization of VGLUT1 & PSD-95 detected no statistical difference between genotypes (F = 2.820, p = 0.0794). (**D**) Representative images from molecular cell layer of dentate gyrus. (**E**) Graphical representation of image location used in synaptic density analysis. (**F**) Synaptic density quantified by colocalization of VGLUT1 & PSD-95 detected statistically significant increase in synaptic density in Q175DN mice compared to wildtype (p = 0.0367) and zQ175 (p = 0.0438) mice. (F = 4.513, p = 0.0217). Error bars denote mean ± SEM. One-way ANOVA with Tukey’s post hoc test. n = 3 animals/genotype, 3 slices/animal. #*p* < 0.1, \**p* < 0.05, \*\**p* < 0.01, \*\*\**p* < 0.001.

VGlut1 is also expressed throughout the hippocampus where also marks excitatory synapses and it is responsible for most of the hippocampal signal conductance [89, 90]. Analyses of synapse density were conducted by colocalizing PSD-95 and VGlut1 in both the CA1 (**Figure 8E-H**) and DG (**Figure 8I-L**). Although we found a trend towards increased PSD-95 puncta number in zQ175 mice compared to WT in the CA1, no significant differences were found in either VGlut1 puncta number of PSD-95/VGlut1 colocalization (**Figure 8F-H**). In the DG, we found no significant differences in either PSD-95, VGlut1, or PSD-95/VGlut1 colocalizations between WT and zQ175 mice. However, contrary to what was expected, we observed a significant increase in synapse density in the DG in Q175DN mice compared to either WT or zQ175 mice (**Figure 8L**). These data suggest that increased aggregation of HTT in the DG of Q175DN not only did not decrease synapse density but rather caused enhanced excitatory signaling.

### Chaperone expression remains largely unchanged in Q175DN mice despite presence of increased mHTT aggregates in both hippocampus and striatum

Deficits in the Heat Shock Response (HSR) have been previously reported in HD cells and mouse models [57, 91] with declining expression of Heat Shock Factor 1 (HSF1) and Heat Shock Proteins (Hsps) and other chaperones in the striatum and other brain regions as disease progresses [57]. Previous reports in other mouse models (R6/2 and HdhQ150) have also shown deficits in Hsp protein expression although RNA levels were not affected [92]. Depletion in HSF1 and Hsps levels are associated with cellular dysfunction and alterations in synapse density. We decided to investigate whether the spare of hippocampal neurons in Q175DN mice was related to changes in the HSR.

We analyzed the levels of HSF1 in the CA1, CA2/3 and DG (granule cell layer and hilus) in WT, zQ175 and Q174DN mice (**Figure 9A, B**). Interestingly, no differences in the expression levels of HSF1 were observed among genotypes throughout the hippocampus except for the CA2/3 where we observed a significant increase in HSF1 levels in Q175DN mice. We also analyzed the expression of two key Hsps, Hsp70 (*Hspa1a*) and Hsp25 (*Hspb1)* using RT-qPCR (**Figure 9C, D**). Hsp expression analyses were conducted from RNA extracted from whole hippocampus. Interestingly, we saw a significant increase in *Hspb1* in zQ175 mice compared to WT, but no significant changes were observed for *Hspa1a* in either zQ175 or Q175DN mice (**Figure 9C, D**). These findings suggest that expression levels of these two Hsps do not necessarily correlate with changes in mHTT aggregation or HSF1 levels, at least in the hippocampus.

**Figure 9.**
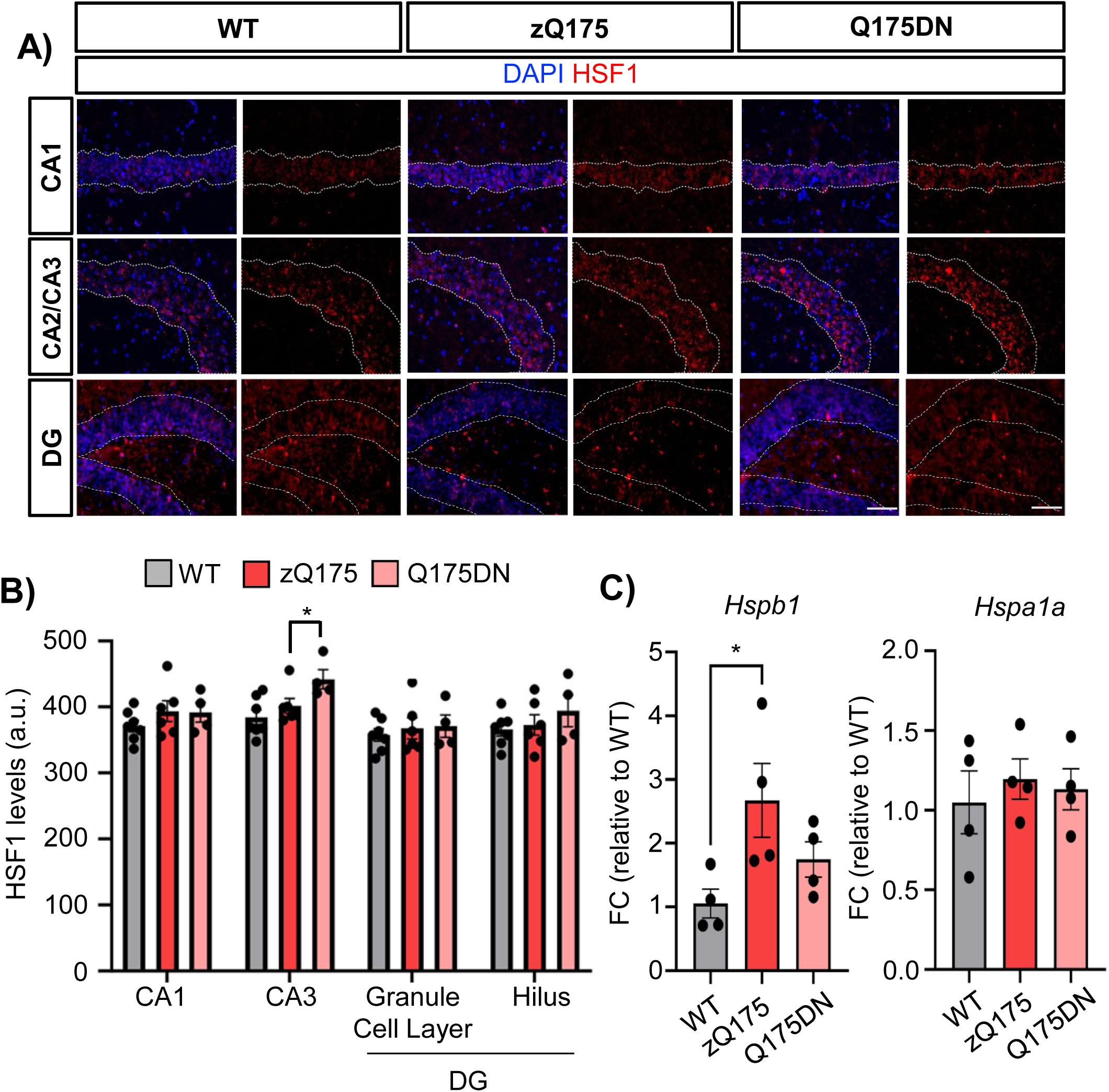
Q175DN mice present higher levels of HSF1 in the CA2/3 region of the hippocampus but no alteration in overall Hsps hippocampal expression. (**A**) Representative immunostainings of HSF1 in the hippocampus of WT (n=7), zQ175 (n=6), and Q175DN (n=4) mice. DAPI stains nuclei. Scale bar, 50 µm. (**B**) Quantification of HSF1 levels (expressed in arbitrary units) in the CA1, CA2/3, DG, and hilus. (**C-D**) Fold change mRNA expression levels for Hsp25 (Hspb1) (**C**) and Hsp70 (Hspa1a) (**D**) in the hippocampus for WT, zQ175 and Q175DN mice at 12 months of age. Data was normalized using GAPDH as control and relativized to WT levels. (n=4 animals/genotype). Error bars denote mean ± SEM. Two-way ANOVA with Sidak’s multiple comparisons in B. One-way ANOVA with Tukey’s post hoc test in C-D. [(B) Interaction: F = 0.5280, df = 6; striatal subregion: F = 5.146, df = 3; genotype: F = 4.588, df = 2)]. Only p-value <0.05 are shown. \**p* < 0.05.

## DISCUSSION

In this work we have conducted a comparative analysis in the response to the presence of mHTT aggregates between two brain regions with essential functions in the control of motor and cognitive behaviors, the striatum and hippocampus (**Table 1**). We used two different mouse models of HD the zQ175 and in house generated Q175DN mice. Our analyses revealed a differential vulnerability between the striatum and hippocampus to the enhanced presence of mHTT aggregates in Q175DN compared to zQ175 and demonstrated the enhanced spare capacity of hippocampal neurons to mHTT toxicity. These findings have important implications for the understanding of differential cellular toxicity mediated by mHTT and the contribution of these two brain regions to the symptomatology of HD.

Our findings in the in-house generated Q175DN mice support previous reports showing increased HTT aggregation and pathology in the striatum of Q175DN and Q175FDN mice, as well as more pronounced behavioral abnormalities compared to zQ175 [45]. This effect is linked to the derepressive influence of removing the Neo cassette upstream of the m*Htt* allele. We confirmed that total *Htt* expression levels were elevated in Q175DN mice relative to zQ175 in the striatum. Notably, since the expression of the endogenous *Htt* allele remains stable in zQ175 [65], any increase in total *Htt* expression in Q175DN can be attributed to the heightened expression of the mutant allele. While mHTT aggregates are typically viewed as a hallmark of HD, some studies have questioned their role in toxicity and cell death susceptibility [93]. Additionally, various mouse models of HD have shown no clear correlation between mHTT inclusion and pathogenic phenotype [94, 95] pointing to mutant HTT expression, solubility, or HTT exon1 splicing as more direct factors in driving pathology. However, the direct relationship between *Htt* transcript levels and HTT aggregate load observed between zQ175 and Q175DN in the striatum makes it difficult to disentangle the influence of each in the pathological assessments we conducted. Nevertheless, although the enhanced pathological changes in Q175DN may not stem from HTT aggregation alone, we utilized this parameter to evaluate the association between HTT aggregation load, vulnerability of different brain subregions, and neuropathological assessments and found key correlations of these parameters in the striatum but not in the hippocampus.

The dorsal striatum has been widely documented to present enhanced degeneration compared to other striatal regions in HD in both mice and humans [6, 58, 96, 97]. Interestingly, when we evaluated the number or size of aggregates across the different regions of the striatum for both zQ175 and Q175DN mice, we found no difference in the number or size of mHTT aggregates between the dorsal regions (dm and dl) and more central regions (cm and cl). However, we found a direct correlation between DARPP-32 levels and EM48 throughout the striatum and specifically in the dm. The increased vulnerability reported for the dorsal striatum might be attributed to a combination of several other factors including the higher density of medium spiny neurons (MSNs) in this region, the specific expression patterns of proteins like Rhes, which moves and transport mHTT between neurons and brain regions [98, 99], and the increased and complex innervation of this region by the corticostriatal and thalamostriatal pathways[83], leading to a greater susceptibility to the toxic effects of mHTT.

In addition to enhanced mHTT aggregation throughout the striatum, we also found a significant increase in the accumulation of mHTT aggregates in all tested regions of the hippocampus (CA1, CA2/3, DG) in the Q175DN compared to the zQ175 with the greatest differential accumulation in the CA2/3 and DG. Interestingly, our EM48 data reported little aggregation in the CA1 of zQ175 mice at 12 months and almost nonexistent EM48^+^ aggregates in the CA2/3 and DG suggesting that the zQ175 hippocampus remains spared from mHTT aggregates until very late in disease. However, studies in this same mouse model using alternative antibodies targeting HTTexon 1 and using formic acid to disrupt HTT amyloid structure have been able to detect the presence of HTT aggregates in the hippocampus as early as 2 months of age [30]. These results indicate that the hippocampus is subject to mHTT aggregation as any other brain region but differences in detection between different HTT antibodies suggest the structure and/or composition of those aggregates might differ compared to the striatum. As such, we found an enhanced size of HTT aggregates in the hippocampus of Q175DN compared to zQ175 mice, but not in the striatum. Interestingly, the levels of HTT aggregation in the hippocampus of zQ175 and Q175DN mice did not correlate with changes in total *Htt* transcript. These results suggest that phenomena other than increased mutant *Htt* expression might be responsible for the enhanced aggregation in Q175DN compared to zQ175 in this brain region. A possible explanation could be the presence of differential posttranslational modifications that could affect the stability and/or propensity of the HTT protein to aggregate, even if total transcript levels remain constant.

Despite enhanced EM48 signal throughout the hippocampus in Q175DN mice, we did not observe signs of increased pathology in any of the tested regions when compared to either zQ175 or WT mice. These results are in concordance with previous reports comparing the susceptibility to mHTT-mediated death in striatal and hippocampal neurons when using ectopic expression of a truncated fragment of mouse HTT (first 480 amino acids) containing 68 CAG repeats showing induced cell death in striatal but not in hippocampal neurons [100]. However, there are contradicting reports that question such hippocampal resilience. Studies in HN33 cells, which are immortalized cells originated from the fusion of mouse hippocampal neurons and neuroblastoma cells, showed high sensitivity to full-length human mHTT induced apoptosis [101] and early stereological studies in postmortem hippocampi from HD patients showed hippocampal degeneration in the CA1 [102]. While the use of immunofluorescence analyses and measures of pathology in the brain, like those conducted in this study, are powerful tools to assess differential vulnerabilities to mHTT, they may not be sensitive enough to detect subtle pathological changes. Therefore, we cannot completely rule out the absence of hippocampal pathology in either zQ175 or Q175DN mice. More sophisticated studies using ^11^C-UCB-J small-animal PET imaging of the synaptic vesicle glycoprotein 2A (SV2A), a vesicular protein that regulates the release of both glutamate and GABA, revealed a significant depletion in SV2A in the hippocampus of Q175DN mice, at least at older ages, suggesting hippocampal synaptic dysfunction [52]. We found increased PSD-95/VGlut1 colocalization in the DG of the hippocampus of Q175DN mice, which reflects the density of excitatory synapses that respond to glutamate. It is possible that increase in colocalized PSD-95/VGlut1 puncta may reflect a compensatory mechanism in response to disruptions in other neurotransmitter systems and the imbalance between excitatory and inhibitory signaling, ultimately affecting the overall function of the hippocampus and its contribution to cognitive processes. However, while these studies might suggest a disruption in the hippocampal circuitry, several neuroimaging and other brain analyses in patients have systematically considered the hippocampus a spared region in HD [103–105], supporting the enhanced resistance of hippocampal neurons to mHTT when compared to striatal neurons.

Early studies in various mouse models have demonstrated that regional differences in susceptibility to mHTT cannot be explained by regional differences in Htt expression, as such, the hippocampus has significantly greater levels of endogenous Htt than the striatum [106]. Somatic instability of the expanded CAG repeat has been implicated as a factor mediating the selective striatal neurodegeneration in HD [107, 108]. Studies in mice and humans showed that expanded CAG repeats undergo further expansion-biased somatic instability in a tissue-specific manner, with striatum and cortex displaying the longest repeat lengths, and significantly affected by age[109–111]. Therefore, it is reasonable to hypothesize that the differences in pathology seen between the striatum and hippocampus might be attributed to a difference in CAG instability. Indeed, MiSeq sequencing of brain DNA samples from HD patients revealed the putamen presented the highest ratio of CAG somatic expansions (10.3%) immediately followed by the hippocampus (6.6%) and the lowest ratio found in the cerebellum (4.8%)[112]. Unfortunately, a comprehensive comparison of CAG instability across brain regions and HD mouse models (transgenic and knock-in HD mice) is lacking. However, those studies assessing CAG instability in transgenic HD mice (R6/1 and R6/2) in which the hippocampus was included showed CAG instability did not differ between striatum and hippocampus [107]. These results imply that, at least in mouse models, the differences in susceptibility between striatum and hippocampus may not be necessarily explained by differences in CAG instability.

Our results reopen the debate of the actual toxicity of mHTT aggregates and put into question what mechanisms drive this differential vulnerability between striatum and other brain regions [93, 113]. We and others have previously shown that HD striatal cells are unable to activate a proper protein stress response and suffer from the abnormal degradation of the stress protective transcription factor HSF1 [57, 91, 114]. Depletion of HSF1 in the striatum is accompanied by downregulation of several chaperones, genes involved in protein quality control mechanisms and other synaptic genes regulated by HSF1 [57]. However, instead of signs of pathology in the hippocampus of Q175DN we found enhanced synaptic density in the DG, decreased number of Iba1^+^ microglia in the CA1 and CA3, and increased HSF1 levels in the CA3 compared to zQ175 mice which might imply the activation of an efficient stress response in Q175DN mice that protects the hippocampus from mHTT accumulation.

The question is how these results explain HD symptomatology. Cognitive deficits including alterations in recognition and spatial learning and memory have been widely documented in HD patients and reproduced in several HD mouse models [36, 41, 115, 116]. Traditionally, the hippocampus and entorhinal cortex have been considered the brain regions primarily involved in these behaviors [27, 28]. Studies in Q175FDN mice showed deficits in acquisition and reversal learning when tested in the object location task (OLT) [45], a cognitive test that primarily evaluates spatial learning[117]. Both Q175FDN and Q175DN mice have also shown exacerbated deficits in recognition memory compared to zQ175 when tested on the novel object recognition (NOR) test [45, 118], a highly validated test to assess non-spatial learning of object identity [119]. However, the lack of signs of overt hippocampal pathology in either the Q175DN or zQ175 compared to WT mice suggests that brain regions other than the hippocampus might play a role in the cognitive behaviors employed to assess cognitive deficits in HD mice. This is supported by lesion studies in the rat CA1 showing no performance effect in the OLT [120] while lesion studies in the dorsomedial striatum of mice have exhibited delayed spatial learning when tested in other cognitive tests [121]. Several recent studies have demonstrated that tasks such as the OLT, NOR or other behavioral assessments of learning and memory are also highly influenced by the striatum [25, 26, 122]. Interestingly, recent evidence has indicated an interaction between the hippocampal and striatal systems in spatial memory [23], with the right-posterior hippocampus implicated in memory of boundary-related locations and the right dorsal striatum involved in memory of landmark-related location [26]. This interaction has been examined through functional MRI in control and HD patients showing a dynamic and compensatory interaction between these two brain regions in the regulation of some cognitive behaviors [123]. Taking this evidence into consideration, it is possible that striatum degeneration would be the major driver of the cognitive deficits seen in HD mice and patients.

In summary, our data provides further support for the selective vulnerability of neuronal subtypes to mHTT toxicity that contributes to explaining specific behavioral alterations in HD. In the presence of enhanced hippocampal mHTT aggregation, we observed limited pathological markers of hippocampal neurodegeneration; from these findings, we conclude that cognitive deficits in HD may not be primarily caused by mHTT-mediated degeneration of the hippocampus. Additional insight into the Q175DN model’s cognitive phenotypes tested through hippocampal-independent behavioral assays may allow for an improved understanding of the contribution of mHTT and striatal degeneration to cognitive symptoms characteristic of HD. We therefore encourage additional investigations into alternative mechanisms driving cognitive symptoms associated with HD. We also suggest that the enhanced spare capacity observed in hippocampal neurons of Q175DN mice to mHTT aggregation demonstrates the value of this mouse model as a tool to understand the fundamental susceptibility differences to mHTT toxicity between different neuronal subtypes.

## Acknowledgements

Figures 1A, S1A, and S1C were created using BioRender.com. M.A.S. is supported by grants from the University of Minnesota’s Undergraduate Research Opportunities Program.

## Data Availability Statement

Data sharing is not applicable to this article as no datasets were generated or analyzed during this study.

## Supplementary Material

Includes nine supplementary figures (1-9).

## Supplementary Figure legends

**Supplementary Figure 1. Genotyping scheme of neomycin cassette excision in Q175DN mice. (A)** Schematic representation of primer binding for insertion of expanded *HTT* Exon 1. **(B)** Representative agarose gel image of WT, zQ175, and Q175DN amplicons using HdhI, HdhII, HdhIII and HdhIV primers. **(C)** Schematic representation of primer binding for excision of neomycin cassette. **(D)** Representative agarose gel image of WT, zQ175, and Q175DN amplicons using DNeoI and DNeoII primers.

**Supplementary Figure 2. Comparative analysis between EM48 immunofluorescence and DAB staining**. (A) Representative EM48 DAB stainings and immunofluorescence in the cm striatum of zQ175 and Q175DN mice. (B) Data was presented as Fold change EM48 aggregate detections relative to zQ175 (n=1 animals/genotype). (C) Representative EM48 DAB stainings and immunofluorescence in the CA1 of zQ175 and Q175DN mice. (D) Data presented as Fold change EM48 aggregate detections relative to zQ175 (n=1 animals/genotype). Scale bar, 90 µm.

**Supplementary Figure 3. Comparative analysis in the ability to detect HTT aggregates between EM48 and other HTT antibodies**. (A) Graphical representation of HTT antibodies and their antigen. (B) Representative EM48, rbHtt, 1C2, and MAB2166 DAB stainings of WT, zQ175, and Q175DN mice. Scale bar, 90 µm.

**Supplementary Figure 4. Comparative analyses for EM48 puncta across different regions of the hippocampus**. Hippocampal mHTT aggregation quantified by number of EM48^+^ puncta per mm^2^ in Q175DN and zQ175 mice across the CA1, CA2/3, and DG (n=6 zQ175, n=4 Q175DN). Error bars denote mean ± SEM. Two-way ANOVA with Sidak’s multiple comparisons. Interaction: F = 5.916, df = 2; hippocampal region: F = 15.72, df = 2; genotype: F = 34.99, df = 1.

**Supplementary Figure 5. Total *Htt* transcript levels in the striatum and hippocampus**. qPCR analyses for total HTT in ∼12 mths old WT, zQ175, and Q175DN in the striatum (A) and hippocampus (B). Data was normalized using GAPDH as control gene and presented as Fold change mRNA expression levels for mouse *Htt* relative to WT (n=4 animals/genotype). Error bars denote mean ± SEM. One-way ANOVA with Tukey’s post hoc test. \**p* < 0.05, \*\**p* <0.01.

**Supplementary Figure 6. No difference in weight in Q175DN mice compared to zQ175 mice.** Quantification of mouse weight at end of life was conducted in two separate cohorts in 12M and 18M WT, Q175DN, and zQ175 mice. Error bars denote mean ± SEM. One-way ANOVA with Tukey’s post hoc test. (12M) Interaction: F=4.965, p=0.0221; (18M) F=100.5, p<0.0001).

**Supplementary Figure 7. Darpp32 signal is differentially altered throughout the different striatum subregions in Q175DN mice and correlations with EM48 levels**. A) Quantification of mean fluorescence intensity (expressed in arbitrary units) in the dm, dl, cm, and cl striatum of WT (n=9), zQ175 (n=7), and Q175DN (n=4) mice. Error bars denote mean ± SEM. Two-way ANOVA with Tukey’s post-hoc test. Interaction: F = 6.555, df = 6; striatal subregion: F = 27.83, df = 3; genotype: F = 217.0, df = 2. B-E) Simple linear regression analyses between number of EM48 puncta per mm^2^ in the dm (B, F=5.373, dF=8), dl (C, F=3.464, dF=8), cm (D, F=25.29, dF=8), and cl (E, F=4.866, dF=8) striatum.

**Supplementary Figure 8. Comparative analysis of the granule cell layer thickness of the different regions of the hippocampus**. Quantification of granule cell layer thickness via NeuN immunofluorescence staining in the hippocampus of WT (n=4), zQ175 (n=4), and Q175DN (n=4). Comparative analysis was performed using images obtained from experiments in figure 4. Error bars denote mean ± SEM. Two-way ANOVA with Tukey’s post-hoc test. Interaction: F = .9061, dF = 3; hippocampal subregion: F = 14.11, df = 8.

**Supplementary Figure 9. Number of GFAP+ astrocytes is higher in the Hippocampus compared to the Striatum across all genotypes**. Quantification of GFAP^+^ cells per 0.34 mm^2^ in the hippocampus and striatum of WT (n=7), zQ175 (n=6), and Q175DN (n=4). Number of cells are calculated as the average of counts obtained in the CA1, CA2/3, and DG hippocampal regions or the cl, cm, dl and dm striatum. Error bars denote mean ± SEM. One-way ANOVA with Tukey’s post-hoc test, F= 26.08, dF = 32.

