## Supplementary Figures 1-9 for "Enhanced Hippocampal Spare Capacity in Q175DN Mice Despite Elevated mHTT Aggregation"

### Supplementary Material

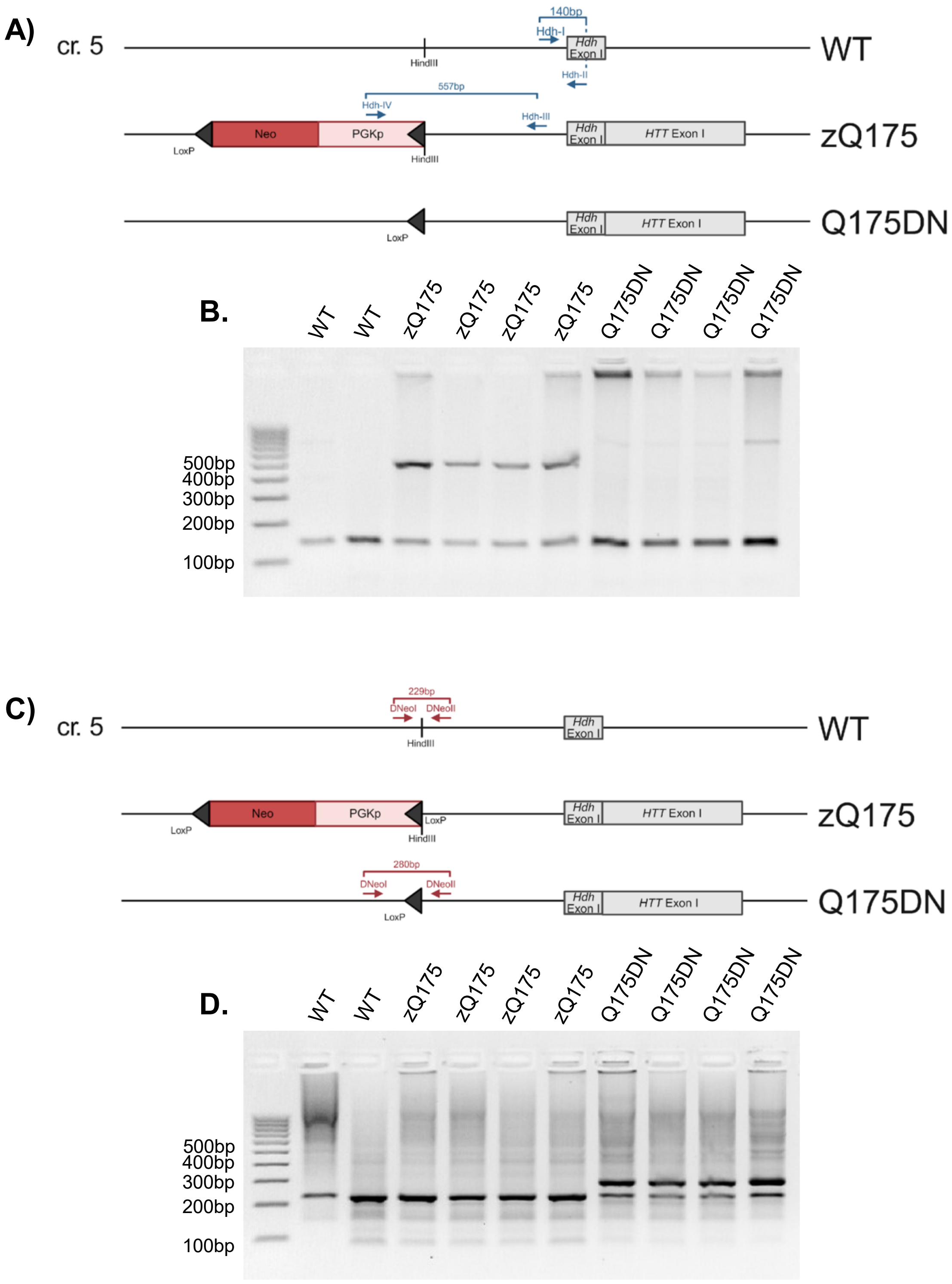


**Supplementary Figure 1**

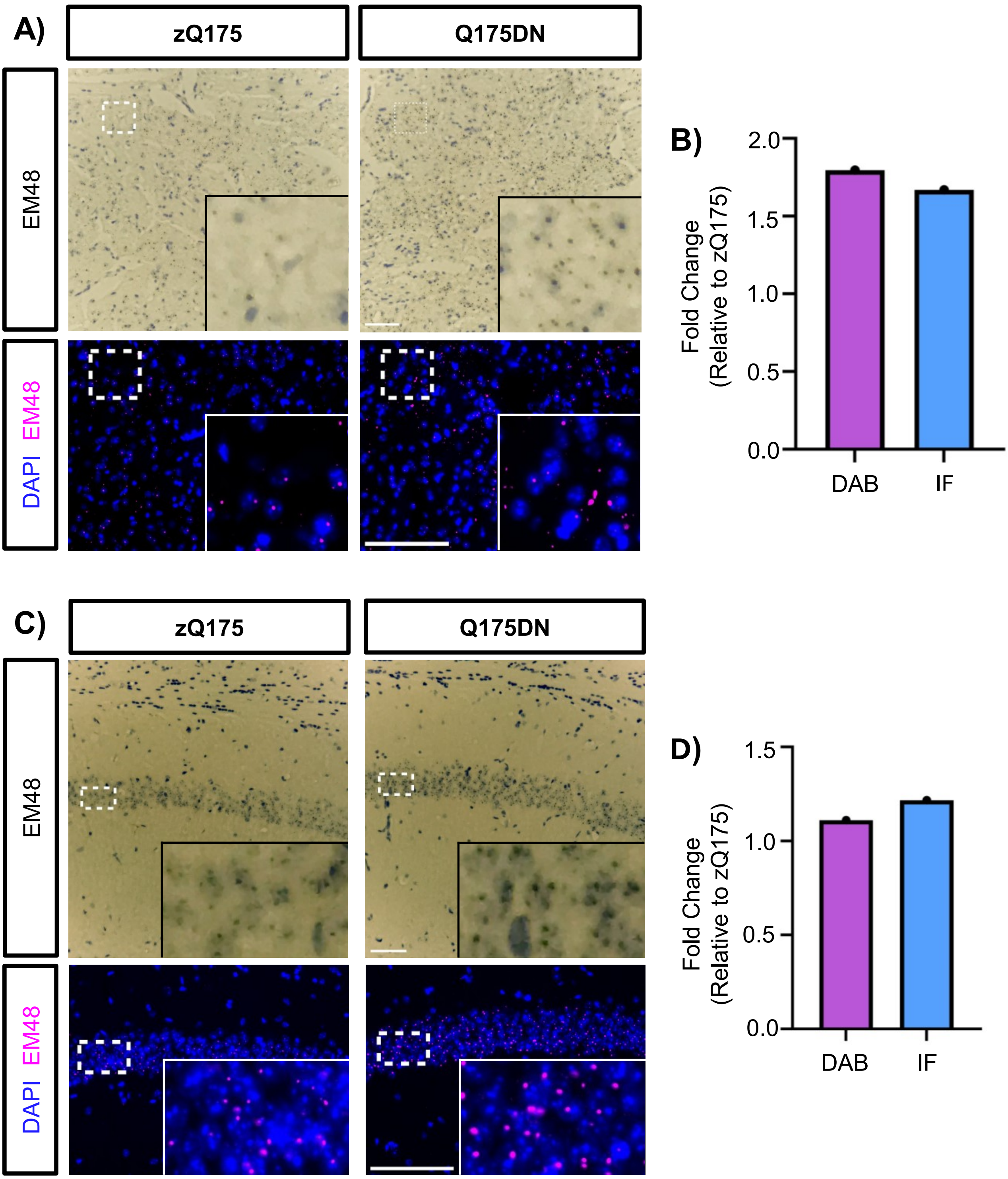


**Supplementary Figure 2**

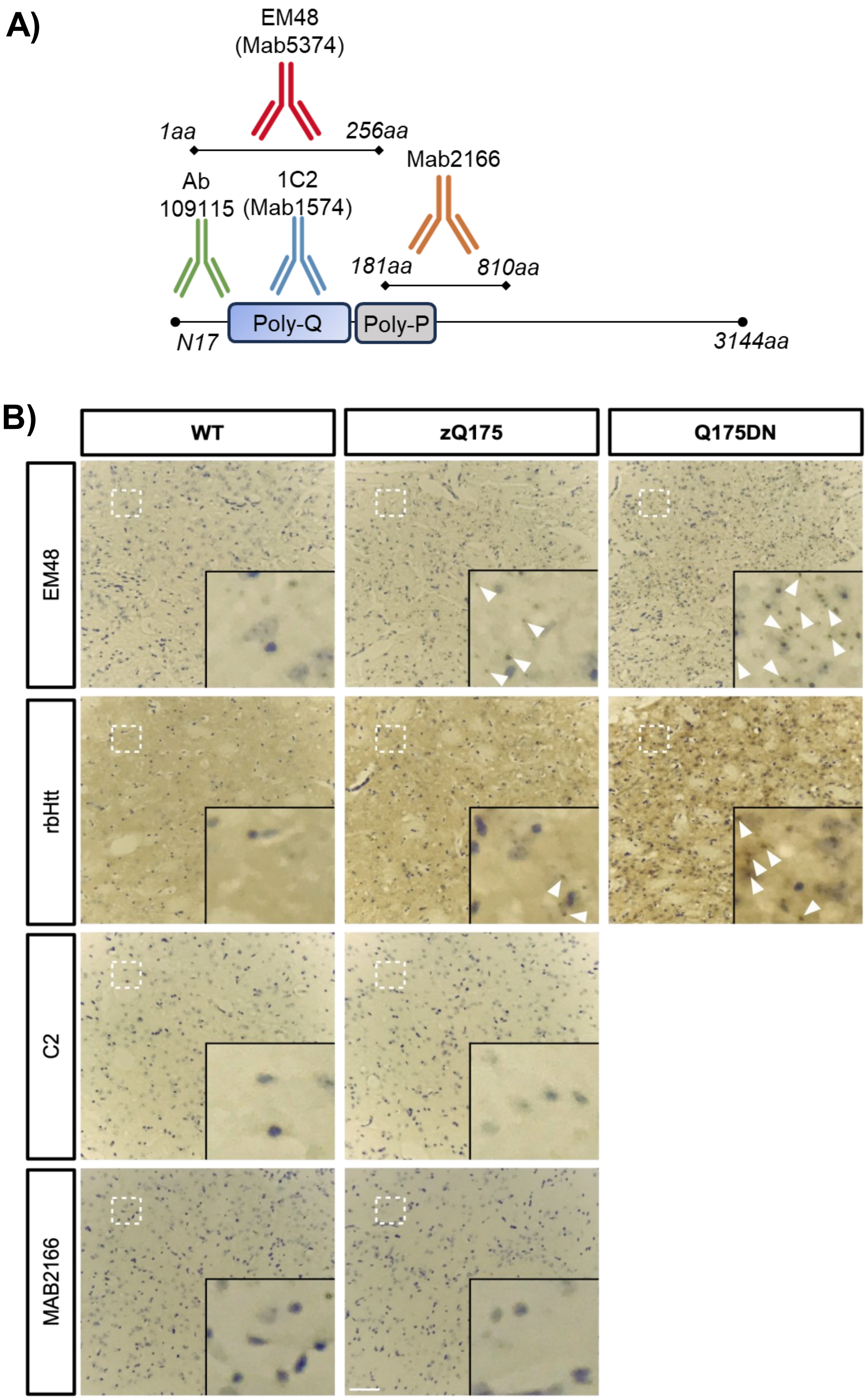


**Supplementary Figure 3**

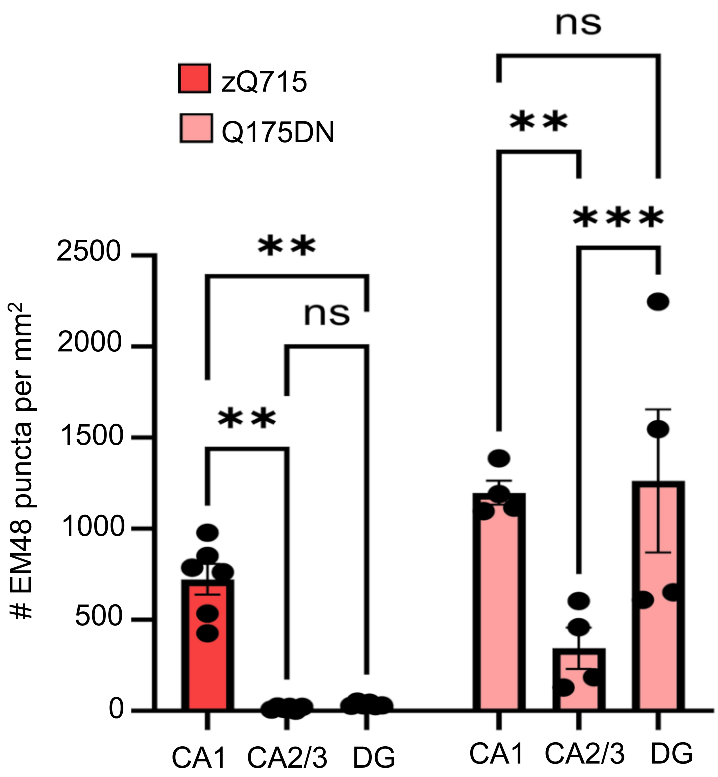


**Supplementary Figure 4**

Total *Htt* transcript levels

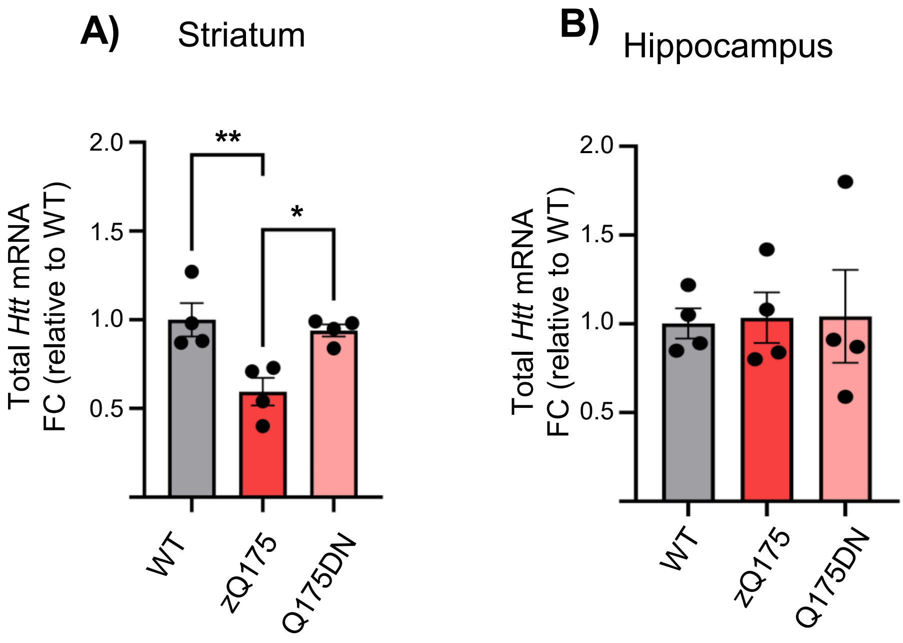


**Supplementary Figure 5**

Body weight

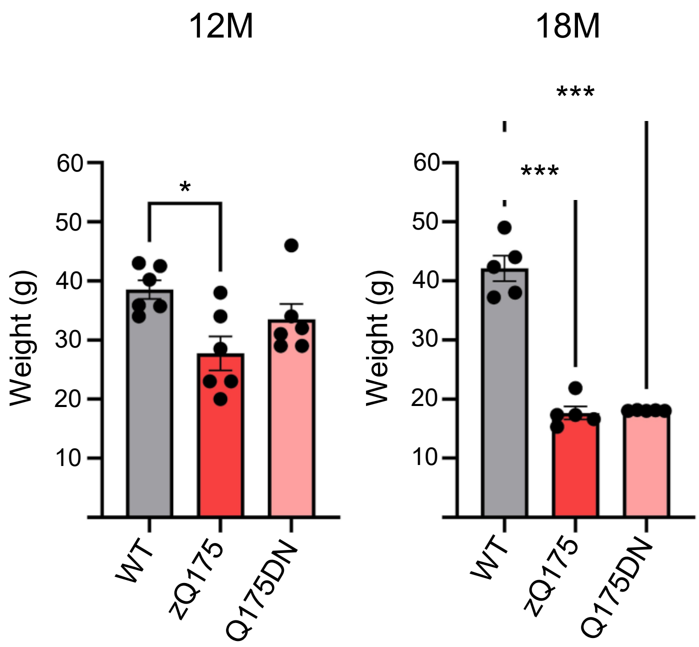


**Supplementary Figure 6**

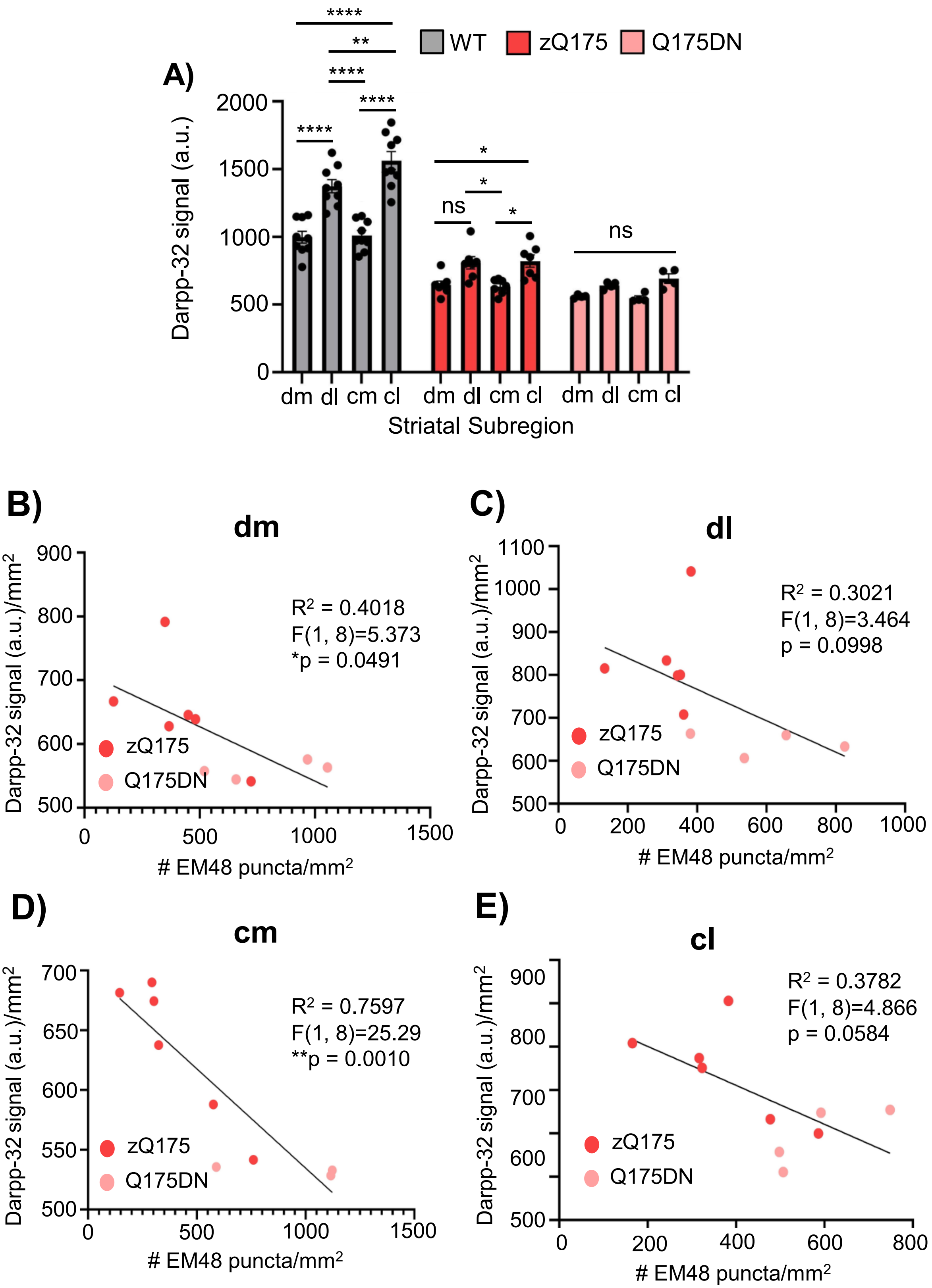


**Supplementary Figure 7**

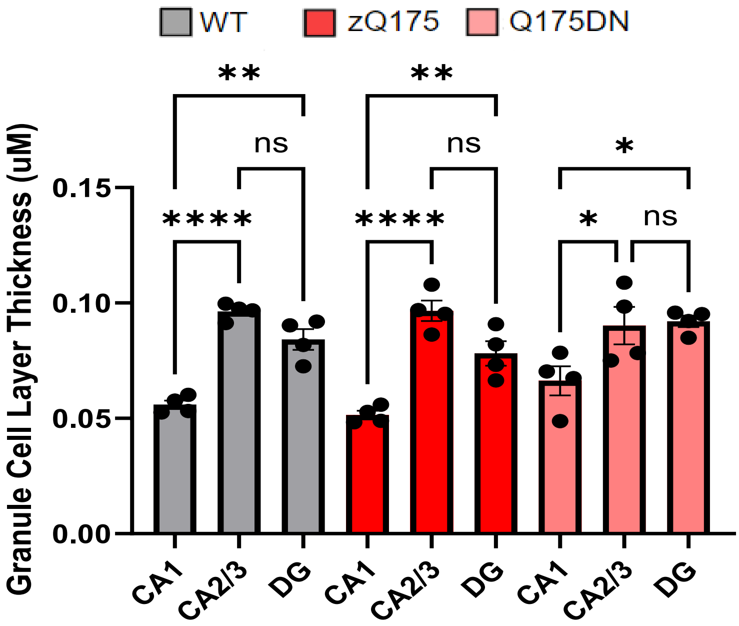


**Supplementary Figure 8**

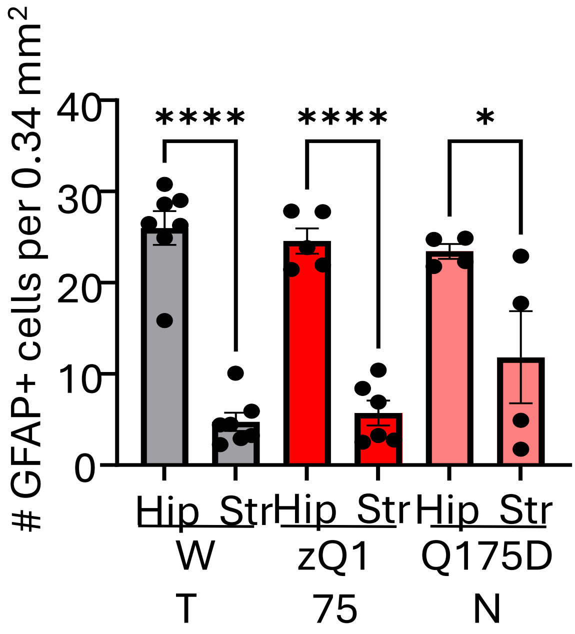


**Supplementary Figure 9**
